# “*Select and retrieve via direct upsampling*” network (SARDU-Net): a data-driven, model-free, deep learning approach for quantitative MRI protocol design

**DOI:** 10.1101/2020.05.26.116491

**Authors:** Francesco Grussu, Stefano B. Blumberg, Marco Battiston, Lebina S. Kakkar, Hongxiang Lin, Andrada Ianuş, Torben Schneider, Saurabh Singh, Roger Bourne, Shonit Punwani, David Atkinson, Claudia A. M. Gandini Wheeler-Kingshott, Eleftheria Panagiotaki, Thomy Mertzanidou, Daniel C. Alexander

## Abstract

**Purpose:** We introduce *“Select and retrieve via direct upsampling”* network (SARDU-Net), a data-driven framework for model-free quantitative MRI (qMRI) protocol design, and demonstrate it on *in vivo* brain and prostate diffusion-relaxation imaging (DRI).

**Methods:** SARDU-Net selects subsets of informative measurements within lengthy pilot scans, without the requirement to identify tissue parameters for which to optimise for. The algorithm consists of a *selector*, identifying measurement subsets, and a *predictor*, estimating fully-sampled signals from the subsets. We implement both using deep neural networks, which are trained jointly end-to-end. We demonstrate the algorithm on brain (32 diffusion-/T1-weightings) and prostate (16 diffusion-/T2-weightings) DRI scans acquired on 3 healthy volunteers on two separate 3T Philips systems each. We used SARDU-Net to identify sub-protocols of fixed size, assessing the reproducibility of the procedure and testing sub-protocols for their potential to inform multi-contrast analyses via *T1-weighted spherical mean diffusion tensor* (T1-SMDT, brain) and *hybrid multi-dimensional* MRI (HM-MRI, prostate) modelling.

**Results:** In both brain and prostate, SARDU-Net identifies sub-protocols that maximise information content in a reproducible manner across training instantiations. The sub-protocols enable multi-contrast modelling for which they were not optimised explicitly, providing robust T1-SMDT and HM-MRI maps and goodness-of-fit in the top 5% against extensive sub-protocol comparisons.

**Conclusions:** SARDU-Net gives new opportunities to identify economical but informative qMRI protocols from a subset of the pilot scans that can be used for acquisition-time-sensitive applications. The simple architecture makes the algorithm easy to train when exhaustive searches are intractable, and applicable to a variety of anatomical contexts.

## 1. Introduction

Quantitative MRI (qMRI) techniques enable the estimation of biophysical properties of imaged tissues from multi-contrast images (1), providing promising system-independent biomarkers in several clinical contexts. Notable examples include: relaxation times, useful to assess myelination in the brain (2) or luminal structures in the prostate (3); diffusion characteristics, linked to cytoarchitecture in various anatomical districts (4–9); blood flow (10); mechanical stiffness of organs such as liver (11); tissue temperature (12).

qMRI can potentially overcome the key limitations of routine clinical MRI, e.g. its limited sensitivity and specificity to early and diffuse alterations that often precede the appearance of focal lesions (13,14). Importantly, recent advances in acquisition have increased dramatically the number of images that can be acquired per unit time (15–17), enabling rich multi-modal qMRI sampling schemes, such as joint diffusion-relaxation imaging (DRI) (18–20). Such novel approaches exploit complementary information from multiple MRI contrasts, and may enable better estimation of microstructural properties compared to single-contrast methods (21,22). Nonetheless, the increased acquisition complexity makes the design of clinically viable protocols challenging. Clinical acquisitions should capture salient signal features within vast sampling spaces in acceptable times, thereby trading off between scan duration and information content.

Previous literature has dealt extensively with qMRI protocol optimisation, i.e. with the design of informative samplings given a specified scan time. Examples include the design of optimal diffusion-weighting protocols (4,5,23–26); number and spacing of temporal sampling in relaxometry (27,28); DRI sampling (18). These previous studies adopt different optimisation strategies, such as Cramér-Rao lower bound (CRLB) minimisation based on Fisher information (29), Monte Carlo (MC) samplings (30), mutual information computation (31), empirical approaches (32) or discrete searches (33). Importantly, these previous optimisation approaches rely on fixed, *a priori* representation of measured signals, such as biophysical models (e.g. multi-compartment models (4,7,9,34)) or phenomenological descriptors (e.g. cumulant expansions (35) or continuous distributions of signal sources (18)). Such representations are indeed useful to capture salient patterns in response to changes in the prescribed MRI pulse sequence. However, adopting *a priori* representations inevitably leads to strong assumptions. Models are often formulated with the healthy tissue in mind (4), which determines the number and properties of water compartments (4,9) included in the models. Moreover, users may need to choose sets of tissue parameter values for which to perform the experiment-design optimisation (29), which typically only consider thermal noise (36) as a source of signal variability, ignoring instrument-dependent factors (37) and physiological noise (38). Ultimately this may limit the generalisability of optimised protocols in real clinical settings and in presence of complex pathophysiological processes.

Here we introduce the *“Select and retrieve via direct upsampling”* network (SARDU-Net), a new data-driven framework for qMRI protocol design that does not rely on any *a priori* explicit signal model or representation. SARDU-Net selects a subset of qMRI measurements that best enables the estimation of a large and comprehensively-sampled qMRI signals. Here we demonstrate it by concatenating two deep neural networks (DNNs), which are trained on real-world *in vivo* qMRI measurements end-to-end. Following previous preliminary investigation (39), we present the implementation of SARDU-Net and demonstrate its capability of finding economical but informative sub-protocols within lengthy state-of-the-art qMRI scans, namely joint DRI of the brain (17) and prostate (40–42), whose deployment in clinical settings is subject to high time pressure.

## 2. Methods

Below we introduce our algorithm and describe experiments performed on data acquired in healthy volunteers in ethically-approved sessions after obtaining informed written consent.

### 2.1. SARDU-Net algorithm

We consider the problem of identifying *D* informative qMRI measurements within an input data set of *in vivo* voxel signals, each made of unidimensional arrays of *M* > *D* measurements. To this end, we couple a *selector*, which extracts a subset of size *D*, and a *predictor*, which estimates fully-sampled M-measurement sets from the subsets (Figure 1). We optimise the two jointly under the hypothesis that the most informative subset enables the best reconstruction of fully-sampled signals. We demonstrate our selector-predictor framework implementing these two as fully-connected DNNs, inspired by deep autoencoders (43,44), recent unsupervised learning tools that hold promise to guide MRI sampling in vast acquisition spaces (45). The selector DNN takes a voxel signal made of *M* measurements *s* = [*S*_1_… *s*_M_]^T^ as input, and outputs a set of *M* corresponding scores **w** = [*w*_1_… *w*_M_]^T^, of which *M* – *D* are zero. Afterwards, the Hadamard product

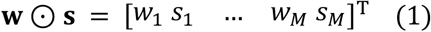

is evaluated to select a subset of *D* out of *M* measurements, and passed to the predictor DNN. The predictor outputs **u** = [*u*_1_… *u*_M_]^T^, an estimate of the fully-sampled signal s derived directly from the subset **w**⊙**s**.

**FIGURE 1.**
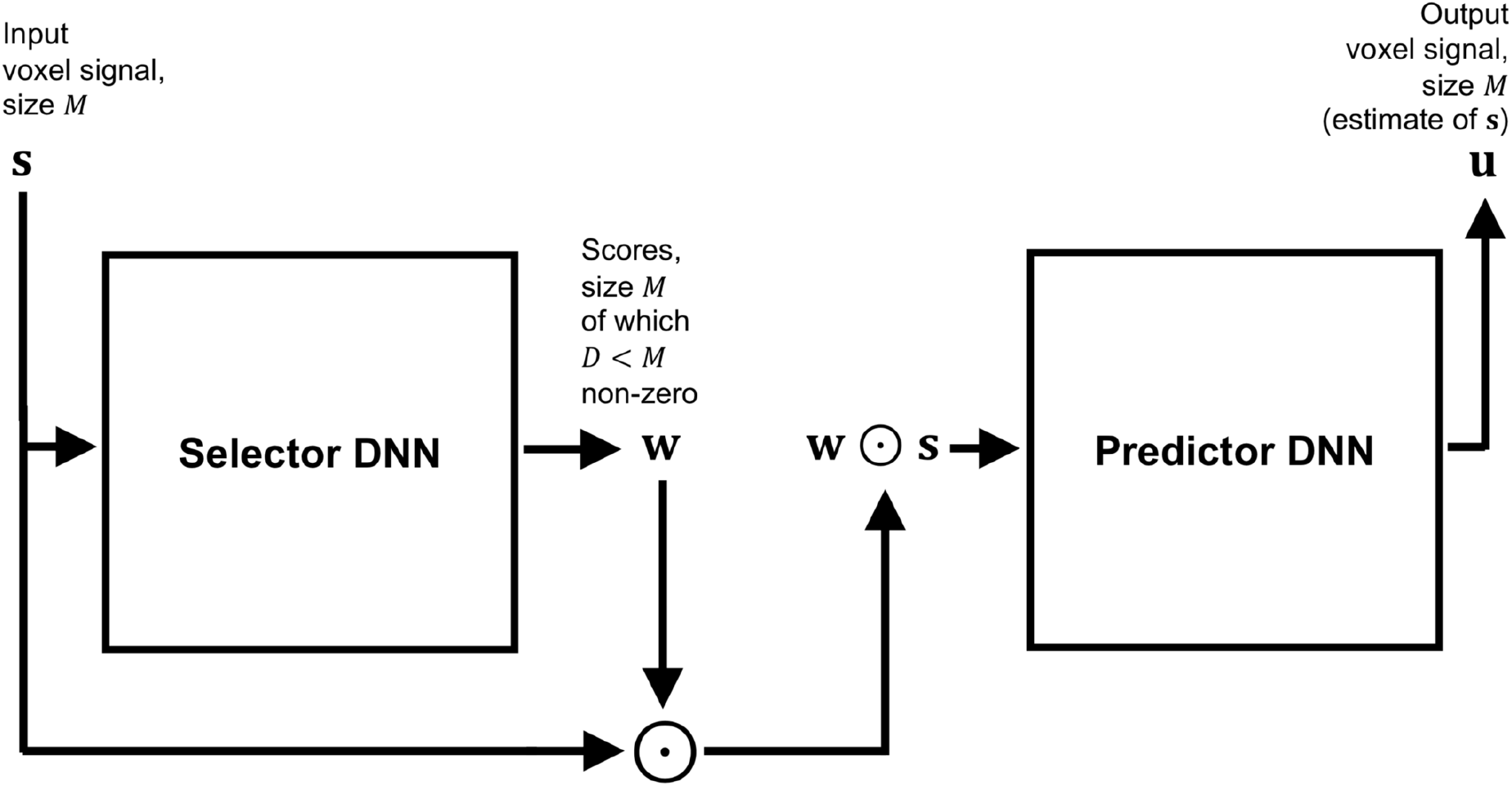
Block diagram summarising the architecture of SARDU-Net, made of two fully-connected DNN that are optimised jointly end-to-end to find optimal subsets of measurements within lengthy quantitative MRI acquisitions. The first DNN is a *selector*: it takes as input a signal made of multiple MRI measurements from a voxel, and outputs a corresponding set of scores, which nullify non-salient measurements effectively extracting a subset of the fully-sampled signal. The second DNN is a *predictor*: it aims to retrieve the fully-sampled signal from the subset received from the selector.

Both selector and predictor are constructed as multi-layer, fully-connected feedforward DNNs, where each layer is obtained as a linear matrix operation followed by element-wise Rectified Linear Units (ReLU). Additionally, the activations of the selector output neurons are normalised to add up to 1 via *softmin* normalisation, and then thresholded so that only the *D* top-firing neurons are kept. The remaining *M* – *D* neurons are zeroed, so that the output scores w effectively select *D* measurements. The selector and predictor are optimised jointly end-to-end to find a subset of measurements that carries the most information about the fully-sampled signal. For this, a loss function

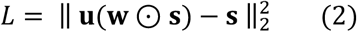

measuring the ℓ^2^-norm of the reconstruction error (i.e. mean squared error (MSE)) is minimised via back-propagation (46) with ADAM optimisation (47) and dropout regularisation (48). In practice, the differentiable product **w**⊙**s** enables the propagation of error gradients from the predictor to the selector, and hence their joint optimisation.

For optimisation, input voxels intensities are normalised as

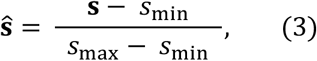

where *s*_min_ = 10^-6^ and *s*_max_ is the 99th percentile of the range of variation of the signal intensity across the whole input set. Afterwards, voxels are split at random into actual training and validation sets. Training voxels are grouped and passed through the network in minibatches; during one such forward network pass, the selector’s output neuron activations are averaged over the mini-batch before computing **w**, and network parameters then updated. The whole set of training voxels is the input to the system for a fixed number of epochs, during which the loss function is evaluated on both training and validation set. In the latter case, the selector DNN is temporarily deactivated and the scores **w** are fixed to the latest values provided by training. Ultimately, a trained SARDU-Net is obtained by deploying the parameters at the epoch with minimum validation loss.

### 2.2. Brain MRI

#### 2.2.1. Acquisition

We performed brain DRI scans on 3 healthy volunteers (2 females, 1 male) using a 3T Philips Ingenia CX system. DRI featured joint diffusion-/T1-weightings, achieved by varying diffusion weighting and inversion time TI (17). A multi-slice saturation inversion recovery (SIR) (49) DW EPI sequence was used, with the vendor’s 32-channel head coil for reception. Salient sequence parameters were: 48 axial-oblique slices, 2.4 mm-thick; field-of-view: 230×230 mm^2^; in-plane resolution: 2.4×2.4 mm^2^; repetition time TR = 2563 ms; TE = 90 ms; saturation delay TS = 300 ms; SENSE factor: 2; multiband factor: 3; readout bandwidth: 2.51 KHz/pixel. Scans were performed with 32 unique (*b*,TI) values among *b* = {0, 1000, 2000, 3000} s/mm^2^ × TI = {70, 320, 570, 820, 1070, 1320, 1570, 1820} ms. For each (*b*,TI) pair, 3 images were acquired for *b* = 0 and 21 isotropic-distributed gradient directions for non-zero *b*-values, optimising directional distribution across the 3 *b*-shells according to (50). This corresponded to 528 EPI images in total, with scan time of 45min:12sec (MRI parameters in Supporting Information Tables S1). Additionally, one *b* = 0 image with reversed phase encoding direction was acquired for distortion correction.

#### 2.2.2. Post-processing

Brain scans were denoised with Marchenko-Pastur Principal Component Analysis (MP-PCA) (51) (kernel: 5×5×5 voxels) and noise floor mitigated with a custom-written Matlab (The MathWorks, Inc., Natick, Massachusetts, USA) implementation of the method of moments (52). Afterwards, motion and eddy current distortions were mitigated via affine co-registration based on NiftyReg (http://cmictig.cs.ucl.ac.uk/wiki/index.php/NiftyReg), with each volume co-registered with reg_aladin to the mean of all 528 EPI images. Finally, FSL topup (53) and bet (54) were used to mitigate EPI distortions and segment the brain. The median signal across all voxels and measurements was used to re-scale image intensities prior to downstream processing.

#### 2.2.3. Experiments

We studied the ability of SARDU-Net to select informative sub-protocols within the set of DRI measurements. We followed a leave-one-out approach and used two out of three subjects to train a SARDU-Net in turn. The remaining subject was then used to test whether SARDU-Net selected an informative sub-protocol. For this demonstration we focussed for simplicity on (*b*,TI) sub-protocols, and did not consider gradient direction dependence, as in related literature (29). We fed SARDU-Net with directionally-averaged DW signals (55) at fixed (*b*,TI), which are commonly referred to as *powder-averaged* or *spherical mean* signals. Directional averaging provides measurements that are not confounded by the underlying fibre orientation distribution (56), and is a common step in several DW MRI techniques (57–59).

##### SARDU-Net training

Sub-protocols of *D* = {16,8,4} out of *M* = 32 measurements were searched, training a SARDU-Net for 300 epochs (80% and 20% of voxels as training/validation sets; four hidden layers for selector DNN with {32,20,24,28,32} neurons for *D* = 16, {32,14,20,26,32} for *D* = 8, {32,11,18,25,32} for *D* = 4; architecture of the predictor mirroring that of the selector; 18 different sets of learning options within mini-batch size = {100,800,1500} voxels × learning rate = {10^-4^,10^-3^} × dropout regularisation = {0.0,0.2,0.4}). Training was performed 8 times for each learning option configuration, initialising the DNNs randomly each time to assess reproducibility.

##### Multi-contrast analysis

We adapted a previous approach (60), which modelled brain white matter inversion recovery DW measurements at b-value *b*, gradient direction **g**, inversion time TI as

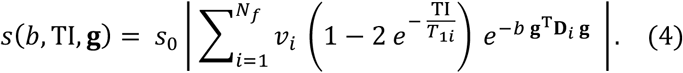

Above *V_i_*, *T*_1*i*_ and **D**_*i*_ are the volume fraction, longitudinal relaxation time and cylindrically-symmetric diffusion tensor of fibre population *i* = 1,…, *N_f_*. Here we adapted Equation 4 by i) including the effect of the saturation pulse (fixed saturation-inversion delay, TS = 300ms); ii) considering directionally-averaged signals; iii) setting *N_f_*= 1 to deploy the model across the whole parenchyma, obtaining

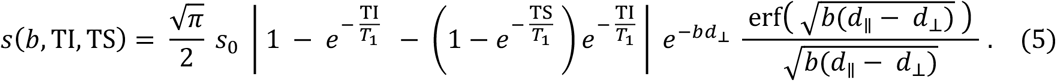

We refer to Equation 5 as *T1-weighted spherical mean diffusion tensor* (T1-SMDT) model, with tissue parameters: *s*_0_ (apparent proton density), *d*_∥_ (fibre parallel diffusivity), *d*_⊥_ (fibre perpendicular diffusivity), *T*_1_ (relaxation time). *d*_∥_ and *d*_⊥_ map properties that are independent of the underlying fibre orientation distribution, and a per-fibre anisotropy index (AI*f*) can be derived (56) as

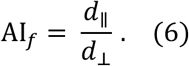

We fitted the T1-SMDT model to the full set of (*b*,TI) measurements using a recent DNN-based fitting approach (61), as available in the qMRI-Net toolbox (62) (link: http://github.com/fragrussu/qMRINet; details in Supporting Information Table S2). For comparison, we repeated T1-SMDT fitting for SARDU-Net sub-protocols as well as for naïve sub-protocols obtained by uniform downsampling of the (*b*,TI) space.

Finally, we performed an extensive numerical evaluation in Matlab to assess SARDU-Net sub-protocols for their ability to inform downstream multi-contrast analyses for which they were not explicitly optimised. We compared SARDU-Net and naïve uniform sub-protocols against 300 random unique sub-protocols of the same size in a dictionary-based model fitting experiment. We generated ~400,000 synthetic signals for each SARDU-Net, naïve uniform and random sub-protocols of the same size by varying tissue parameters (*s*_0_, *d*_∥_, *d*_⊥_, *T*_1_) of Equation 4 within a uniform grid. Afterwards, we obtained the combination of tissue parameters providing the lowest signal MSE, and used it to synthesise fully-sampled signals in each voxel. These were compared to the actual fully-sampled measured signals, obtaining voxel-wise MSEs which were averaged within one brain slice passing through the corpus callosum and containing all tissue types.

### 2.3. Prostate MRI

#### 2.3.1. Acquisition

We acquired *in vivo* DRI scans on 3 healthy males as part of an ongoing study (63) using a 3T Philips Achieva system. DRI featured different diffusion-/T2-weightings, achieved by varying *b*-value and echo time TE. A multi-slice diffusion-weighted (DW) echo planar imaging (EPI) sequence was used, with the vendor’s 32-channel cardiac coil for reception. Salient parameters were: 14 axial slices, 5 mm-thick; field-of-view of 220×220 mm^2^; in-plane resolution of 1.75×1.75 mm^2^; repetition time TR = 2800 ms; SENSE factor: 1.6; half-scan factor: 0.62; 2 averages; 1 coronal REST slab for spatial saturation; readout bandwidth: 2.39 KHz/pixel. Scans were performed with 16 unique (*b*,TE) values among *b* = {0, 500, 1000, 1500} s/mm^2^ × TE = {55, 87, 121, 150} ms. For each (*b*,TE), 3 images were acquired, using 3 orthogonal diffusion gradients when *b* was not zero, for a total of 48 DRI images (total scan time of 6min:15sec; MRI parameters in Supporting Information Table S3).

#### 2.3.2. Post-processing

Prostate scans were denoised slice-by-slice with MP-PCA (51) (kernel: 7×7 voxels), and noise floor mitigated with the method of moments (52). Motion and eddy current distortions were mitigated slice-by-slice by co-registering each 2D image to a reference with affine registration. NiftyReg `reg_aladin` was used, and the reference was obtained as the average of all volumes. Finally, the three images at any fixed (*b*,TE) obtaining 16 unique (*b*,TE) volumes, which were normalised by dividing by the median signal across all volumes and voxels within the prostate (same normalisation factor for all volumes). For this, a prostate mask was manually segmented on the mean EPI image calculated after co-registration in `FSLView` (64).

#### 2.3.3. Experiments

Measurement subsets selected by SARDU-Net were tested for their potential of informing downstream tissue parameter estimation, as for example via HM-MRI (40,41). As in to brain DRI experiments, we followed a leave-one-out approach and used two out of three subjects to train a SARDU-Net in turn. The remaining subject was then used to test the SARDU-Net sub-protocol.

##### SARDU-Net training

For each leave-one-out fold, measurements from prostate voxels of the two training subjects were extracted and assigned at random to training (80% of voxels) and validation (20% of voxels) sets. Sub-protocols of *D* = {12,9} out of *M* = 16 measurements were searched by training a SARDU-Net for 300 epochs. Four hidden layers were used for selector/predictor DNNs (selector: {16,15,14,13,16} neurons for *D* = 12 and {16,14,13,11,16} for *D* = 9; predictor architecture mirroring selector), and 18 different sets of learning options (mini-batch size ={100,800,1500} voxels × learning rate = {10^-4^,10^-3^} × dropout regularisation = {0.0,0.2,0.4}). Moreover, for each configuration of learning options, training was repeated 8 times using different random network initialisation seeds to assess reproducibility.

##### Multi-contrast analysis

SARDU-Net measurement subsets were evaluated for their potential to inform multi-contrast signal analyses. For this evaluation, we adopted one among several potential methods in the literature, i.e. the HM-MRI (41) model, a multi-exponential approach describing the total prostate signal as the sum of luminal, epithelial and stromal components:

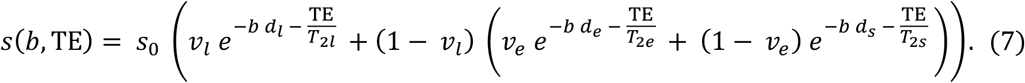

In Equation 7, *s*(*b*,TE) is the prostate signal at a fixed *b*-value and echo time TE. The 9 tissue parameters are: *s*_0_ (apparent proton density); *v_l_* (luminal water voxel volume fraction); *v_e_* (epithelial fraction of non-luminal tissue); *d_l_, d_e_, s_s_* and *T*_2*l*_, *T*_2*e*_, *T*_2*s*_ (ADC and T2 of luminal/epithelial/stromal water).

Firstly, HM-MRI metrics were computed on the fully-sampled scans and on SARDU-Net and on naïve sub-protocols (*D* = {12,9}) for comparison, with naïve sub-protocols obtained by uniform downsampling of the (*b*,TE) measurement space. We used the same DNN-based fitting procedure used for brain DRI (62) and estimated voxel-wise *v_e_, v_e_* and *s*_0_, while fixing compartment-wise ADC and T2 values to literature values (41) (details in Supporting Information Table S4).

Subsequently, we assessed the potential of SARDU-Net sub-protocols to enable downstream analyses for which they were not explicitly optimised for. We performed a similar dictionary-based fitting experiment in Matlab as done for brain DRI (section 2.2.3, *Multicontrast analysis*). In this case, we restricted our analysis to the central slice of each prostate and synthesised a database of ~500,000 reference signals for each 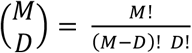 subset of *D* = {12,9} out of *M* = 16 measurements (1,820 for *D* = 12; 11,440 for *D* = 9) by varying tissue parameters uniformly on a grid of previously reported values (41). The combination of parameters providing the lowest sub-protocol MSE was used to synthesise fully-sampled signals and then compute the average mean squared error (MSE) for the fully-sampled protocols within the region-of-interest.

## 3. Results

### 3.1. Brain MRI

Figure 2 shows SARDU-Net selection of *D* = {16,8,4} out of *M* = 32 (*b*,TI) measurements on brain DRI from training fold 1 (complete list of measurement selection available in Supporting Information Tables S5; examples of sub-protocol selection during training available in Supporting Information Figure S6). SARDU-Net sub-protocols sample the full range of *b* and TI values. However, they sample less densely measurements characterised by the lowest SNR levels, as for example strong diffusion-weightings for TI close to the SIR null point.

**FIGURE 2.**
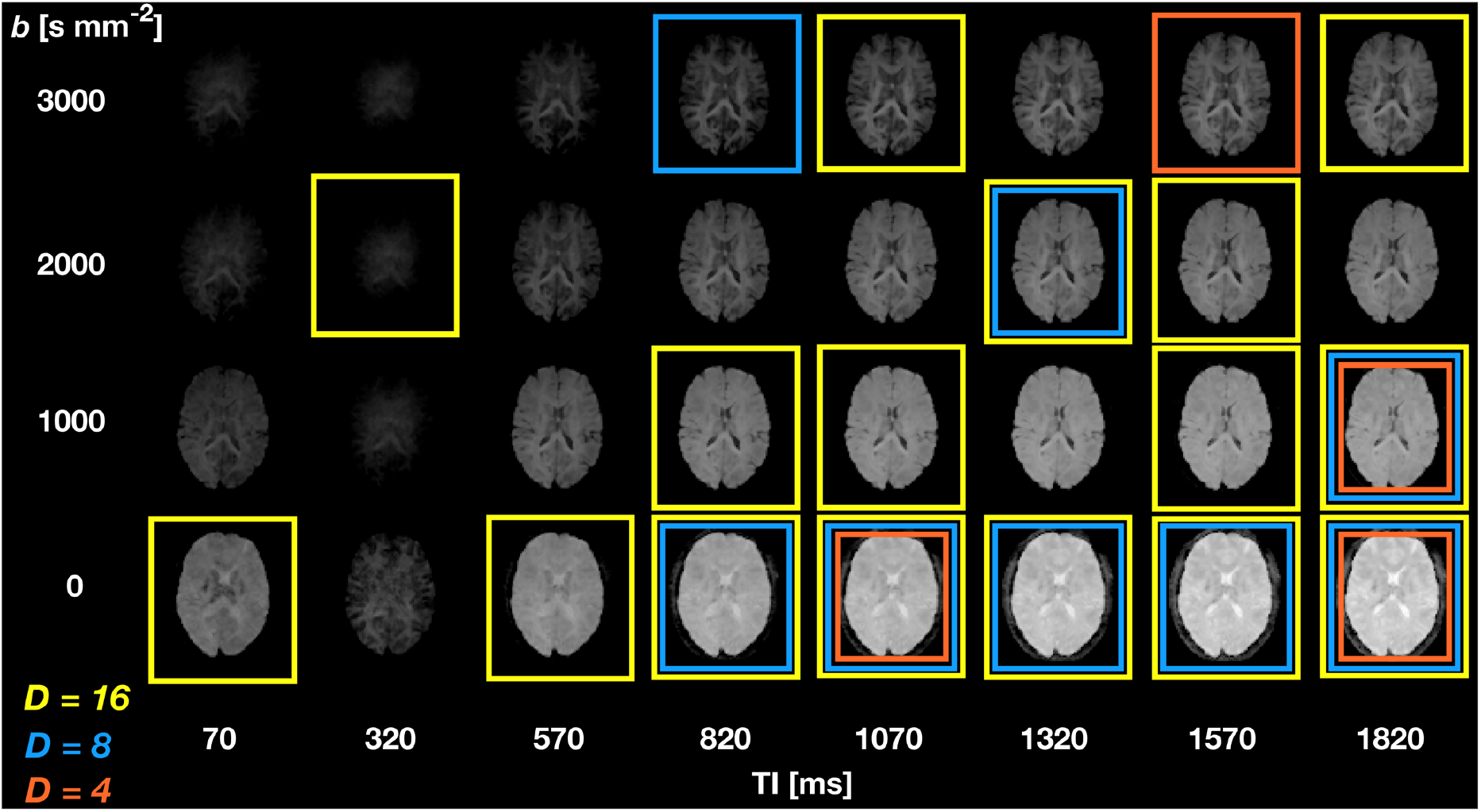
SARDU-Net measurement selection on DRI of the brain. The figure illustrates results from leave-one-out fold 1 (subject 1 left out during training, which is performed on subjects 2 and 3) for selection of *D* = 16 (yellow boxes), and *D* = 8 (blue boxes) and *D* = 4 (orange boxes) out of *M* = 32 (*b*,TI) measurements. The images show spherical mean signals form 1 brain slice of subject 1. Inversion times (TI, delay between inversion pulse and slice excitation) are arranged along different columns, while diffusion-weightings (*b*) along different rows. The saturation delay TS (i.e. delay between saturation pulse and inversion pulse) is fixed for all (*b*,TI) measurements to TS = 300ms.

Figure 3 shows SARDU-Net reproducibility on brain DRI. Results from different subsampling factors are shown in different rows, while results from different leave-one-out training folds are shown in different columns. SARDU-Net measurement selection is consistent across different algorithm initialisations and different training folds. A number of measurements are selected consistently in all cases (e.g. (*b*,TI) = (0 s mm^-2^; 1800 ms)), while other measurements (e.g. (*b*,TI) = (1000 s mm^-2^; 70 ms)) are avoided consistently.

**FIGURE 3.**
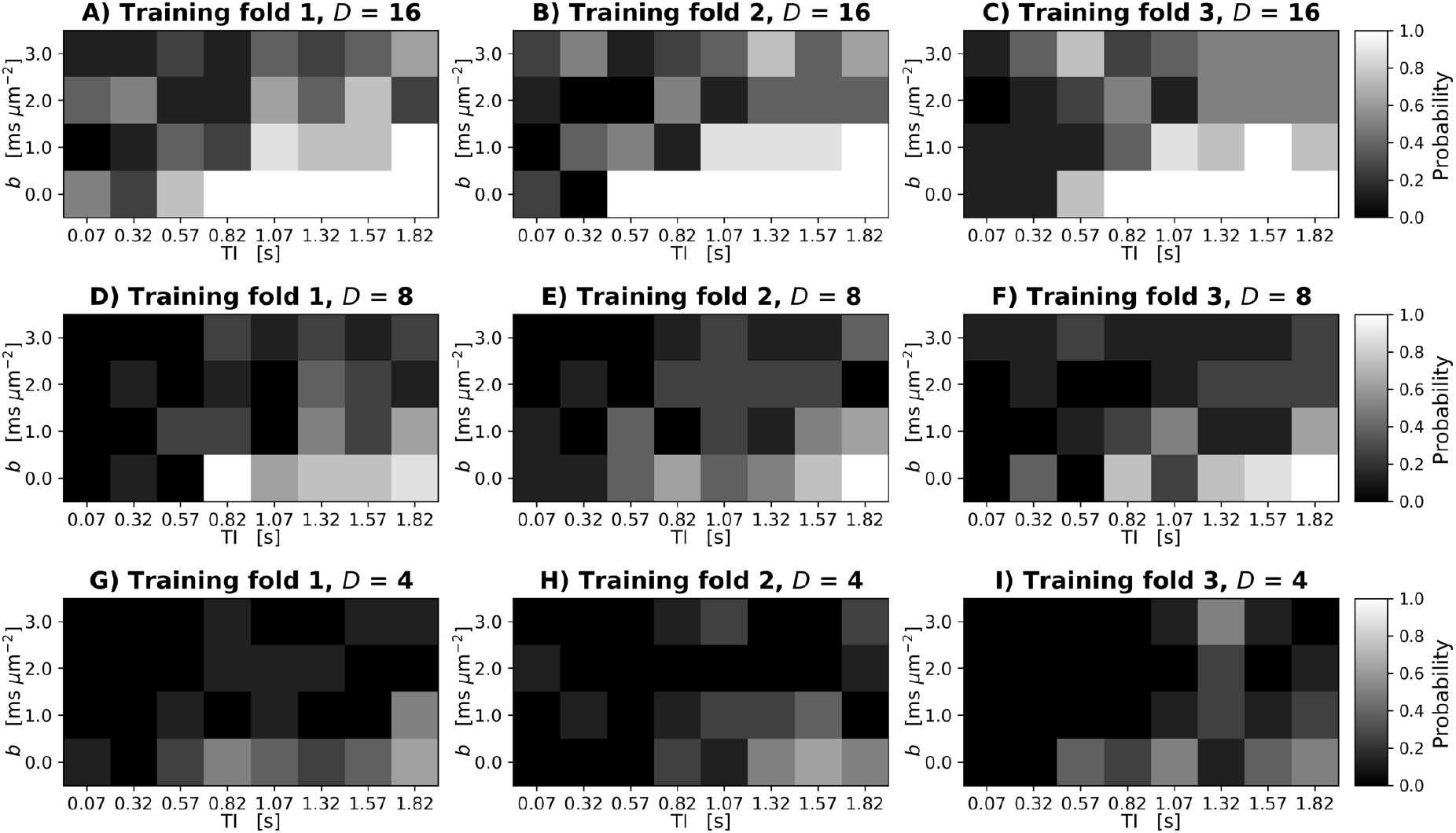
Reproducibility of SARDU-Net measurement selection for *in vivo* brain DRI over different leave-one-out training folds and random initialisations. The normalised 2D histogram in each panel shows the probability of each (*b*,TI) measurement being selected over 8 different repetition of the SARDU-Net training. Each repetition featured a unique random initialisation of the SARDU-Net parameters, with all other training options (i.e. mini-batch size, dropout regularisation, learning rate) fixed to the configuration providing the lowest validation loss. Panels A to C (top row): selection of *D* = 16 out of *M* = 32 measurements (A, left: subject 1 out during training and used for testing; B, middle: subject 2 left out; C, right: subject 3 left out). Panels D to F (middle row): selection of *D* = 8 measurements (D, left: subject 1 out; E, middle: subject 2 out; F, right: subject 3 out). Panels G to I (bottom row): selection of *D* = 4 measurements (G, left: subject 1 out; H, middle: subject 2 out; I, right: subject 3 out).

Figure 4 shows T1-SMDT reference parametric maps from the full protocol as well as those derived from sub-protocols. On visual inspection, parametric maps derived from SARDU-Net and naïve sub-protocols are comparable to the reference when half of the measurements are kept in the sub-protocol (*D* = 16 out of *M* = 32 measurements). However, for stronger downsampling (*D* = 8 and *D* = 4), SARDU-Net sub-protocols preserve key between-tissue contrasts in all parametric maps, unlike naïve sub-protocols.

**FIGURE 4.**
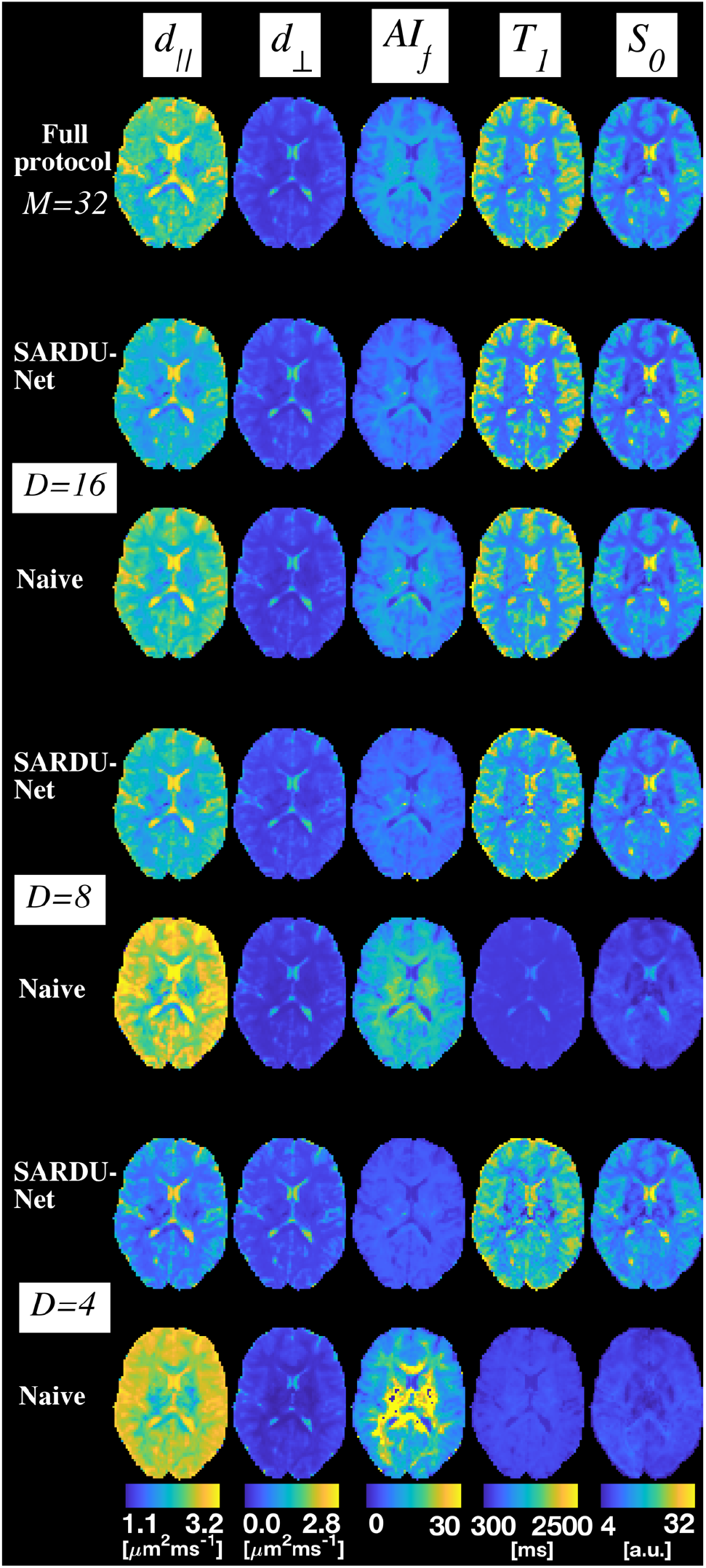
Examples of brain T1-SMDT parametric maps obtained in one subject. Different rows show T1-SMDT metrics from different protocols. From top to bottom: full protocol; SARDU-Net and naïve uniform subprotocol for *D* = 16 measurements; SARDU-Net and naïve uniform subprotocol for *D* = 8 measurements; SARDU-Net and naïve uniform subprotocol for *D* = 4 measurements. Naïve sub-protocols are: (*b*,TI) = {0, 1000, 2000, 300} s mm^-2^ × {70, 570, 1070, 1570} ms for *D* = 16; (*b*,TI) = {0, 1000, 2000, 300} s mm^-2^ × {70, 1070} ms for *D* = 8; (*b*,TI) = {(0, 70), (2000, 70), (1000, 1070), (3000, 1070)} (s mm^-2^, ms) for *D* = 4. From left to right, different T1-SMDT metrics are shown: parallel diffusivity (*d*_∥_); perpendicular diffusivity (*d*_⊥_); anisotropy index (AI_*f*_); longitudinal relaxation time (*T*_1_)); apparent proton density (*s*_0_).

Figure 5 shows MSE distributions obtained when reconstructing fully-sampled brain DRI signals from random sub-protocols with T1-SMDT dictionary fitting. It also reports the MSE values corresponding to SARDU-Net and naïve sub-protocols. SARDU-Net subprotocols exhibit MSE that is in all cases is within the lowest 5%. SARDU-Net and naïve subprotocols have similar MSE when *D* = 16 measurements out *M* = 32 are selected. However, SARDU-Net MSE is considerably lower than naïve sub-protocol MSE for stronger protocol sub-samplings, e.g. *D* = 8 and *D* = 4.

**FIGURE 5.**
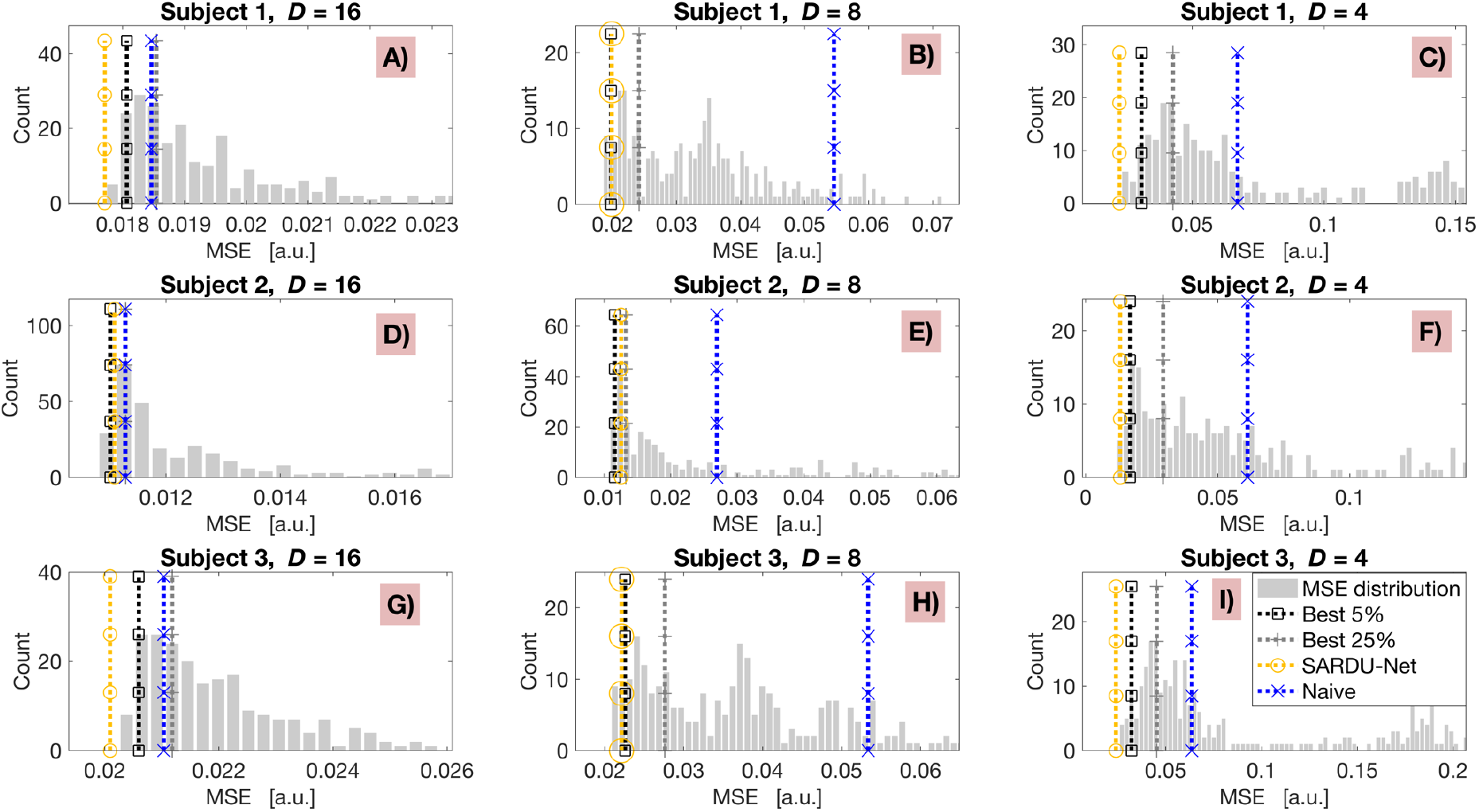
Distribution of MSE from extensive T1-SMDT fitting performed in the test subject of each leave-one-out fold for 300 random sub-protocols (right tail of the distribution cut for visualisation). The MSE for SARDU-Net and naïve sub-protocols are indicated by vertical yellow and blue lines. The top 5% and 25% MSE are flagged by black and dark grey lines. Results obtained on subjects 1, 2 and 3 are reported on top (A, B, C), middle (D, E, F), bottom (G, H, I) panels. Panels A, D, G (left) refer to sub-protocols of size *D* = 16 out of *M* = 32 measurements; panels B, E, H (middle) to sub-protocols of size *D* = 8; panels C, F, I (right) to sub-protocols of size *D* = 4. Naïve sub-protocols are: (*b*,TI) = {0, 1000, 2000, 300} s mm^-2^ × {70, 570, 1070, 1570} ms for *D* = 16; (*b*,TI) = {0, 1000, 2000, 300} s mm^-2^ × {70, 1070} ms for *D* = 8; (*b*,TI) = {(0, 70), (2000, 70), (1000, 1070), (3000, 1070)} (s mm^-2^, ms) for *D* = 4.

### 3.2. Prostate MRI

Figure 6 shows SARDU-Net selection of *D* = 12 and *D* = 9 out of *M* = 16 prostate measurements. The figure refers to training fold 1, with the complete list of measurement selection for all folds in Supporting Information Tables S7. Additionally, an example of the evolution of sub-protocol selection during training is included as Supporting Information Figure S8. Figure 6 demonstrates that SARDU-Net sub-protocols sample the full range of diffusion and relaxation weightings. While measurements with the lowest SNR levels (i.e. maximum *b* and longest TE) are generally not kept in SARDU-Net sub-protocols, in some cases measurements that are selected by SARDU-Net feature lower SNR than measurements that are not selected (e.g. for *D* = 9, (*b*,TE) = (1500 s mm^-2^, 87 ms) is kept, while (*b*,TE) = (500 s mm^-2^, 87 ms) is not).

**FIGURE 6.**
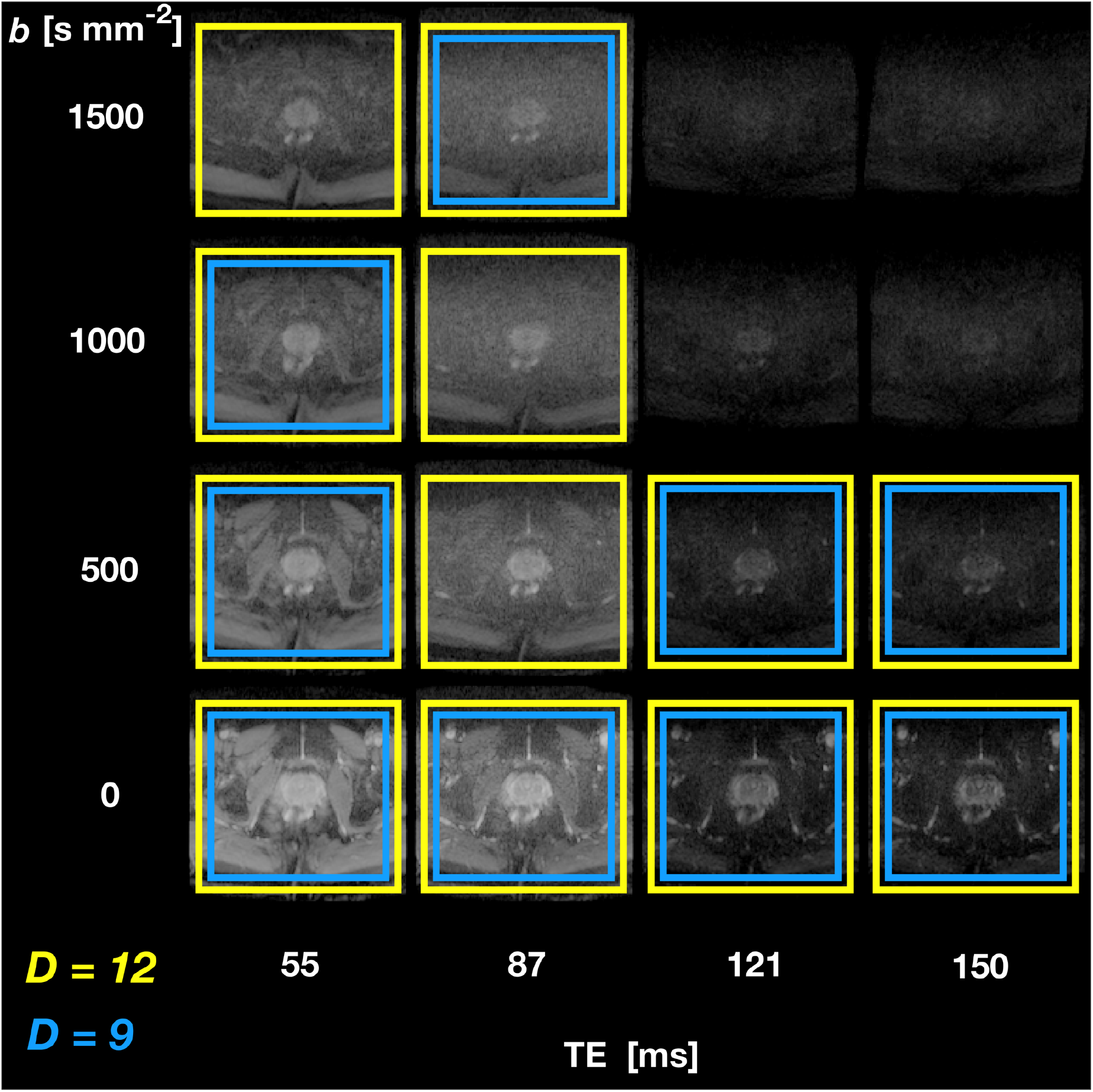
SARDU-Net measurement selection on DRI of the prostate. The figure illustrates results from leave-one-out training fold 1 (subject 1 left out during training, which is performed on subjects 2 and 3) for selection of *D* = 12 (yellow boxes) and *D* = 9 (blue boxes) out of *M* = 16 (*b*,TE) measurements. Echo times (TE) are arranged along different columns, while diffusion-weightings (*b*) along different rows.

Figure 7 shows the reproducibility of SARDU-Net sub-protocol selection across 8 different random initialisations in each training fold. Each panel reports as a 2D histogram the normalised count of each (*b*,TE) measurement being selected over the 8 different random seeds (top row: *D* = 12; bottom row: *D* = 9; different folds along different columns). The illustrations demonstrate that SARDU-Net sub-protocol selection is consistent across leave-one-out folds and random initialisations. For instance, the same 8 measurements out of 12 were consistently selected in all 8 random training repetitions for *D* = 12 in fold 2 and 3, while in no cases SARDU-Net selected the measurement corresponding to sequence parameters (*b*,TE) = (1500 s mm^-2^,150 ms).

**FIGURE 7.**
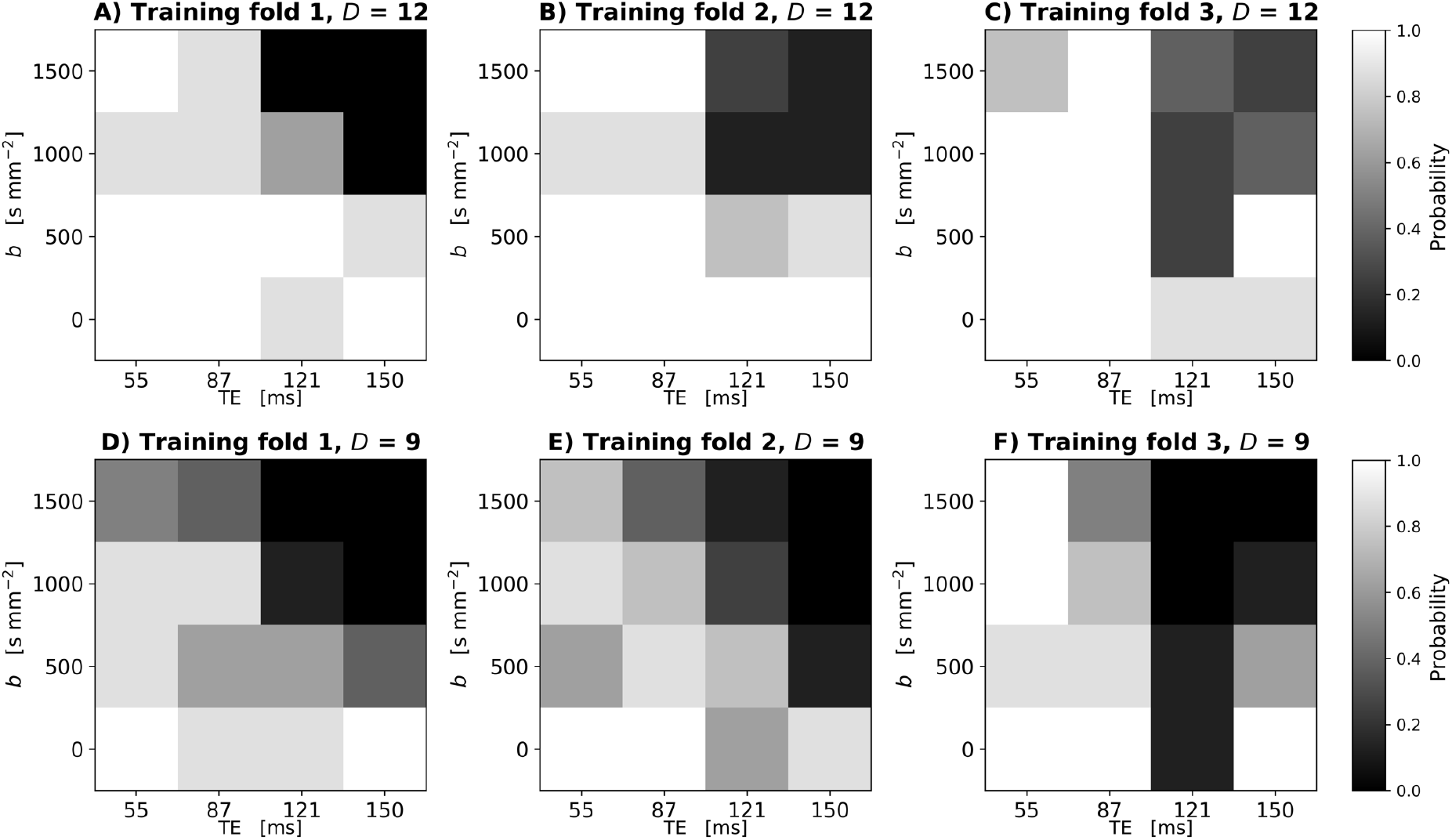
Reproducibility of SARDU-Net measurement selection for *in vivo* prostate DRI over different leave-one-out training folds and random initialisations. The normalised 2D histogram in each panel shows the probability of each (*b*,TE) measurement being selected over 8 different repetition of the SARDU-Net training. Each repetition featured a unique random initialisation of the SARDU-Net parameters, with all other training options (i.e. mini-batch size, dropout regularisation, learning rate) fixed to the configuration providing the lowest validation loss. Panels A to C (top row): selection of *D* = 12 out of *M* = 16 measurements (A, left: subject 1 left out during training and used for testing; B, middle: subject 2 out; C, right: subject 3 out). Panels D to F (bottom row): selection of *D* = 9 measurements (D, left: subject 1 out; E, middle: subject 2 out; F, right: subject 3 out).

Figure 8 illustrates examples of HM-MRI indices obtained on the full protocols as well as on SARDU-Net and naïve sub-protocols. Regional variation of *v*_1_ and *v_e_* maps is in line with known anatomy of the healthy prostate, as for example increased luminal water fraction in the peripheral zone. On visual inspection, metrics obtained from both SARDU-Net and naïve subprotocols show within-prostate contrasts that are qualitatively similar to those obtained from the full protocols. HM-MRI metrics from SARDU-Net sub-protocols appear closer to the fully-sampled reference than naïve sub-protocols, although to a lesser extent than what was observed in Figure 4 for brain DRI.

**FIGURE 8.**
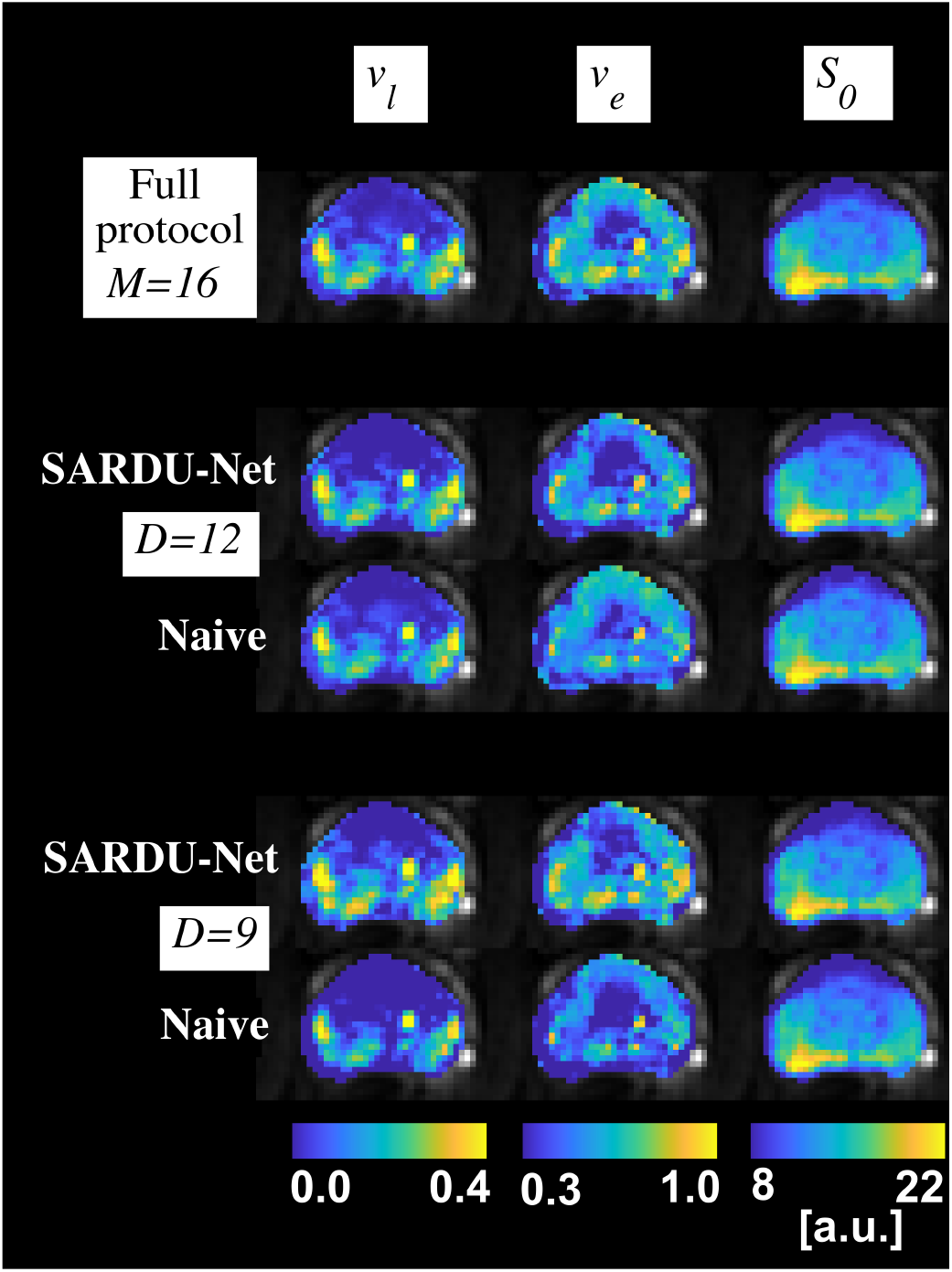
Examples of prostate HM-MRI parametric maps obtained in one subject. Different rows show HM-MRI metrics from different protocols. From top to bottom: full protocol; SARDU-Net and naïve uniform subprotocol for *D* = 12 measurements; SARDU-Net and naïve uniform subprotocol for *D* = 9 measurements. Naïve uniform sub-protocols were: (*b*,TE) = {0, 500, 1000, 1500} s mm^-2^ × {55, 121, 150} ms (D = 12); (*b*,TE) = {0, 1000, 1500} s mm^-2^ × {55, 121, 150} ms (D = 9). From left to right, different HM-MRI metrics are shown: lumen voxel fraction *V*_1_; epithelial tissue fraction *v_e_*; apparent proton density (*s*_0_).

Figure 9 shows MSE distributions obtained by reconstructing fully-sampled prostate DRI signals via HM-MRI dictionary fitting on sub-protocols. Each panel reports the distribution of average MSE obtained for all possible sub-protocols of fixed size on the leave-one-out test subject. Values of MSE obtained for SARDU-Net and naïve sub-protocols are reported explicitly with a vertical bar. SARDU-Net identifies sub-protocol that enable the reconstruction of fully-sampled signal with an error within the lowest 5% in almost all cases. MSE values for SARDU-Net sub-protocols are comparable to naïve sub-protocols for *D* = 12, while they are always considerably lower when subsampling is stronger (*D* = 9).

**FIGURE 9.**
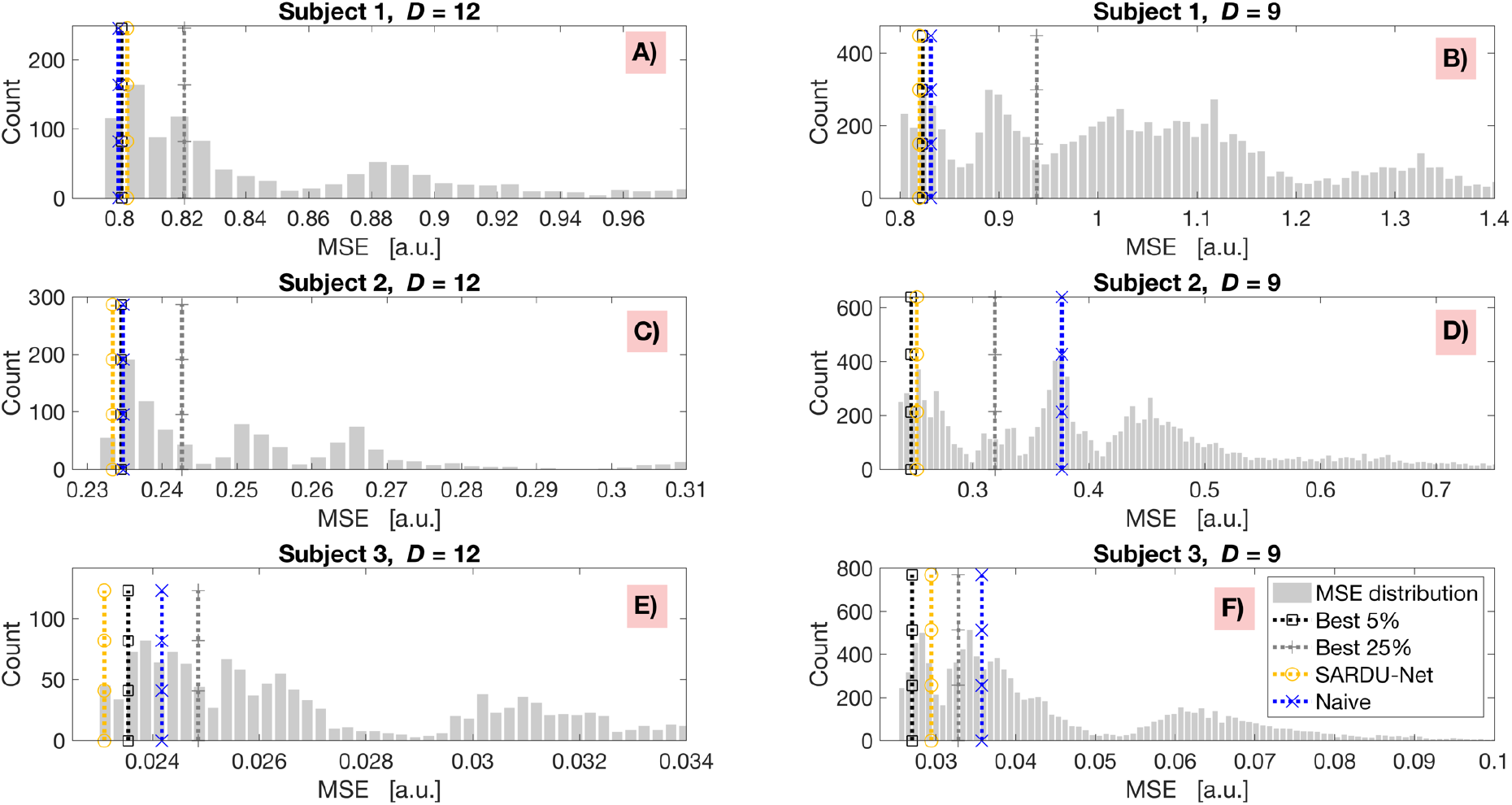
Distribution of MSE from HM-MRI dictionary fitting performed in the test subject of each leave-one-out fold, for all possible sub-protocols of fixed size (right tail of the distribution cut for visualisation). The MSE for SARDU-Net and naïve sub-protocols are indicated by vertical yellow and blue lines. The top 5% and 25% MSE are flagged by black and dark grey lines. Results obtained on subjects 1, 2 and 3 are reported on top (A-B), middle (C-D), bottom (E-F) panels. Panels A, C, E (left) refer to sub-protocols of size *D* = 12 out of *M* = 16 measurements, while panels B, D, F (right) to sub-protocols of size *D* = 9. For *D* = 12, naïve results refer to the protocol with lowest MSE among (*b*,TE) = {0, 500, 1000, 1500} s mm^-2^ × {55, 87, 150} ms and {0, 500, 1500} s mm^-2^ × {55, 87, 121, 150} ms. For *D* = 9, naïve results refer to the protocol with lowest MSE among (*b*,TE) = {0, 500, 1000} s mm^-2^ × {55, 87, 121} ms and {0, 1000, 1500} s mm^-2^ × {87,121,150} ms.

## 4. Discussion

### 4.1. Key findings

This paper presents SARDU-Net, a data-driven, model-free method for qMRI experiment design. SARDU-Net aims to identify informative sub-protocols within lengthy *in vivo* scans to facilitate the deployment of the latest qMRI techniques in contexts where scan time is limited. Our main finding is that SARDU-Net provides a general, robust and reproducible procedure to identify informative sub-protocols. SARDU-Net shows utility across a range of anatomical districts (e.g. brain, prostate) and qMRI contrasts (diffusion, T2, T1), without relying on *a priori* models of the MRI signal.

### 4.2. Sub-protocol selection

We studied DRI scans of the brain and prostate acquired at 3T on two separate groups of healthy volunteers. Brain scans consisted of 32 unique (*b*,TI) measurements via SIR DW imaging, while prostate scans featured 16 unique (*b*,TE) measurements. Data were analysed with SARDU-Net to identify informative sub-protocols within the fully-sampled measurement set, made of 16, 8 and 4 out of 32 for brain and 12 and 9 measurements out of 16 for prostate. The reproducibility of the measurement selection procedure across leave-one-out folds and random initialisations was also assessed.

Our results demonstrate that SARDU-Net identifies informative sub-protocols within densely sampled measurement sets, and that such sub-protocols do not necessarily feature uniform downsampling of the acquisition space. The selected measurements span the whole range of signal weightings for both brain and prostate. Nonetheless, measurements with the lowest SNR levels (i.e. maximum *b* and TI close to neural tissue SIR null point; maximum *b* and longest TE for prostate DRI) are consistently avoided. This suggests that noise is an important factor to consider for qMRI sampling design. While model-based optimisation typically relies on some *a priori* hypotheses on the level and statistics of thermal noise, our fully-data driven approach enables the design of qMRI samplings from real-world SNRs and noise distributions.

Importantly, we characterised the reproducibility of SARDU-Net. Results from both brain and prostate demonstrate that the stability of our sub-protocol selection procedure can enable practical protocol design from a limited number of pilot scans. However, our results also highlight that some variation in sub-protocol selection across training folds ad random algorithm initialisations. The former is likely to originate from intrinsic between-subject variability, and could be minimised by ensuring that the pilot training cohort is large enough to capture biological variability. The latter suggests the presence of different local minima in the algorithm loss function, a known issue in optimisation problems. Such latter variability appears to be small, since a number of key information-carrying measurements are selected consistently. However, in future we aim to reduce the sensitivity of the training procedure to the initial conditions.

### 4.3. Multi-contrast analysis

We tested sub-protocols selected by SARDU-Net for their ability to inform downstream modelbased multi-contrast analyses, for which they were not optimised explicitly. To this end, we adapted a previous brain white matter modelling method to our SIR DWI data (i.e. here referred to as T1-SMDT), and utilised a simple multi-exponential model capturing the joint (*b*,TE) dependence of the MRI signal in the prostate (i.e. the HM-MRI model). Both models were fitted to the full set of measurements and on sub-protocols provided by SARDU-Net, as well as on naïve sub-protocols featuring uniform downsampling of the measurement space. On visual inspection, parametric maps obtained from SARDU-Net are closer to reference maps from full protocols, especially in the brain and when downsampling is stronger, suggesting that SARDU-Net sub-protocols preserve key features of measured signals.

Importantly, parametric maps from both HM-MRI and T1-SMDT models exhibit differences when derived from different DRI protocols. This highlights the intrinsic challenge of inverting highly non-linear models to resolve diffusion/relaxation properties from noisy measurements (8,65). Importantly, we point out that the HM-MRI and T1-SMDT models were simple and convenient choices for our demonstration. Different approaches within a wider landscape of alternative models could have been equally adopted, each with its own advantages and disadvantages. In particular, VERDICT (7) and Relaxed-VERDICT (66) would account for diffusion time in prostate DRI, neglected in this study, while multi-compartment models (8,58,67,68) could be adapted for brain DRI. We reserve such alternative approaches for future investigation.

Finally, we compared SARDU-Net sub-protocols for their ability to capture salient characteristics of input MRI signals against an extensive list of alternative sub-protocols. We used the T1-SMDT and HM-MRI models to reconstruct fully-sampled signals from i) all possible sub-protocols of fixed size for prostate DRI (12 and 9 measurements out of 16) and from ii) 300 random sub-protocols for brain DRI (16, 8 and 4 measurements out of 32), which included naïve sub-protocols featuring uniform measurement space downsampling. Our analyses show that SARDU-Net sub-protocols capture salient features of fully-sampled DRI signals, since in almost all cases they are within the top performing sub-protocols in terms of reconstruction MSE. Importantly, SARDU-Net MSEs are consistently lower than those provided by naïve uniform sub-protocols in both brain and prostate data, with the difference in performance becoming stronger as subsampling becomes more aggressive. This suggests that uniform sampling of DRI measurement spaces is a reasonable choice when a high number of measurements can be taken. However, non-trivial sampling is likely to better capture salient signal characteristics when only a few measurements can be taken.

### 4.4. Methodological considerations

The envisioned deployment of SARDU-Net would require the acquisition of a small number of rich, pilot qMRI scans when a new clinical study is being set up, from which informative subprotocols could be identified given a scan time budget. Pilot scans are typically performed any way for quality control when developing new MRI procedures. Importantly, such pilot scans could be included in subsequent group-level analyses, since the final protocol would be a subset of it.

In this first demonstration, we test our new framework only on healthy volunteers. However, including patients in the pilot training set is imperative to enable the selection of qMRI protocols that capture key signal features typical of pathological tissues. The design of protocols for imaging both health and disease is an issue for our strategy as it is for traditional model-based approaches. We reserve the investigation of measurement selection in disease to future applicative studies. These may exploit data augmentation techniques to increase the number of examples of under-represented pathological signals, as well as from any other tissue whose accurate characterisation may be of interest (e.g. grey matter compared to white matter).

Our approach relies on the hypothesis that the training scans suffice to capture the salient features of input MRI signals, i.e. that they densely sample the qMRI measurement space. The latest acquisition technologies (15) make it achievable to sample two to four sequence parameters (e.g. echo time, inversion time and diffusion encoding) densely in under one hour (17). However, our data-driven approach would become impractical if larger acquisition spaces were of interest, as pilot protocols exceeding the hour would be needed. Related to this point, we acknowledge that our prostate DRI was well under the hour limit (nominal scan time of six minutes), and therefore may not be as representative of rich qMRI samplings as our brain DRI. This was due to the fact that the scan was performed as part of an ongoing MRI study (63). In future, we will explore richer (*b*,TE) samplings and include diffusion time dependence (7,66), which is not considered in this demonstration, to better assess the potential of SARDU-Net for prostate imaging.

Importantly, we remark that our model-free approach is an alternative to previous model-based optimisation strategies (29), which remain valid options when a specific model is the main interest of a study. Here we propose an alternative framework, which could be adopted when multiple downstream analyses are of interest or when the qMRI model is not known to a high degree of confidence at the time of the acquisition, a common situation in several qMRI scenarios.

Another important consideration is that here we compared sub-protocols selected by SARDU-Net to uniform downsamplings of DRI measurement spaces, which were referred to as *naïve* sub-protocols. We acknowledge that alternative *naïve* sub-protocols could have been identified. We avoided naïve sub-protocols that would have provided unfair advantages to SARDU-Net (for instance, all b-values were included in naïve brain DRI sub-protocols even when *D* = 4), and included computational experiments where SARDU-Net sub-protocols are compared to an exhaustive list of alternatives (300 sub-protocols for brain DRI; all subprotocols for prostate DRI). In both cases SARDU-Net sub-protocols enable downstream analyses for which they were not explicitly optimised for, suggesting that our SARDU-Net method does capture the salient features of input MRI signals.

Finally, we showed the utility of coupling and optimising jointly a selector and a predictor. We demonstrated this by implementing both with fully-connected DNNs, given their excellent function approximation properties (69,70). This simple structure suffices to demonstrate the potential and flexibility of data-driven qMRI protocol design, making the algorithm easy to train with limited computational resources when extensive sub-protocols searches become unfeasible (i.e. a situation quickly reached for *M* ~ 30). Nonetheless, we acknowledge that different design choices could be equally valid, as for example genetic searches (33) for the selection stage. In future we will improve the performance and stability of SARDU-Net, for instance by replacing the simple grid search used here to design SARDU-Net architecture and learning options.

### 4.5. Conclusions

SARDU-Net identifies economical but informative qMRI protocols for clinical application under high time pressure. Using a small number of rich pilot acquisitions with long acquisition times, SARDU-Net selects informative sub-protocols given a specific time constraint that capture the salient characteristics of densely-sampled MRI signals.

## Supporting information

Supporting Information Tables S1 and S3 report the full set of MRI parameters. Supporting Information Tables S2 and S4 describe DNN-based model fitting. Supporting Information Tables S5 and S7 report in full SARDU-Net sub-protocols. Supporting Information Figures S6 and S8 show the evolution of SARDU-Net sub-protocols during training.

## Data and code availability statement

Our open-source implementation of SARDU-Net is freely available online (permanent link: http://github.com/fragrussu/sardunet). All analysis scripts written for this paper are also made openly available online (permanent link: http://github.com/fragrussu/PaperScripts/tree/master/sardunet). These depend on the following third-party toolkits:

- FSL: http://fsl.fmrib.ox.ac.uk/fsl/fslwiki;
- NiftyReg: http://cmictig.cs.ucl.ac.uk/wiki/index.php/NiftyReg;
- MP-PCA: http://github.com/NYU-DiffusionMRI/mppca_denoise/blob/master/MPdenoising.m;
- Noise-floor mitigation: https://github.com/fragrussu/MRItools/tree/master/matlabtools/MPio_moments.m;
- Gibbs ringing removal: http://github.com/RafaelNH/gibbs-removal;
- qMRI-Net for DNN-based fitting: http://github.com/fragrussu/qMRINet.

The diffusion gradient direction scheme for the *in vivo* brain data was generated with: http://www.emmanuelcaruyer.com/q-space-sampling.php.

The *in vivo* MRI scans cannot be made openly available online due to privacy issues of clinical data according to GDPR regulations. Researchers interested in accessing the prostate scans can contact Prof Daniel C. Alexander, while researchers interested in the brain scans can contact Prof Claudia A. M. Gandini Wheeler-Kingshott. A data sharing agreement enabling noncommercial research use will be stipulated.

## Acknowledgements

The authors would like to thank the volunteers for their time; the radiographers at UCH and F. Gong for their assistance with prostate MRI protocol set up and data acquisition; Philips Healthcare for assistance in protocol development and for access to research protocols. FG is grateful to R. Tanno, E. Kaden, M. Palombo, P. Slator and C. Jin for useful discussion.

## Funding

This project was funded by the Engineering and Physical Sciences Research Council (EPSRC EP/R006032/1, M020533/1, G007748, I027084, N018702). This project has received funding under the European Union’s Horizon 2020 research and innovation programme under grant agreement No. 634541 and 666992, and from: Rosetrees Trust (UK, funding FG); Prostate Cancer UK Targeted Call 2014 (Translational Research St.2, project reference PG14-018-TR2); Cancer Research UK grant ref. A21099; Spinal Research (UK), Wings for Life (Austria), Craig H. Neilsen Foundation (USA) for jointly funding the INSPIRED study; Wings for Life (#169111); UK Multiple Sclerosis Society (grants 892/08 and 77/2017); the Department of Health’s National Institute for Health Research (NIHR) Biomedical Research Centres and UCLH NIHR Biomedical Research Centre; Champalimaud Centre for the Unknown, Lisbon (Portugal); European Union’s Horizon 2020 research and innovation programme under the Marie Skłodowska-Curie grant agreement No. 101003390. FG is currently supported by the investigator-initiated PREdICT study at the Vall d’Hebron Institute of Oncology (Barcelona), funded by AstraZeneca and CRIS Cancer Foundation.

## Conflicts of interest disclosures

FG is supported by PREdICT, a study co-funded by AstraZeneca in Spain. TS is an employee of Deep Spin (Germany) and previously worked for Philips (UK).

---

**Supporting Information Table S1.**
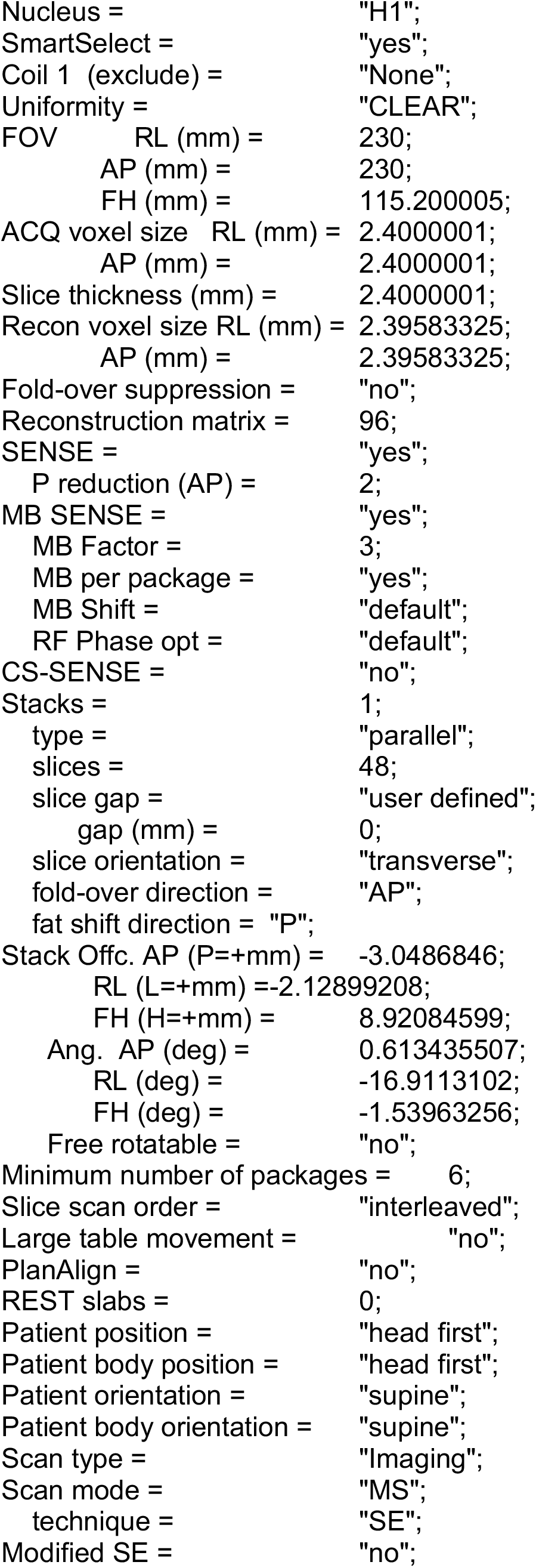
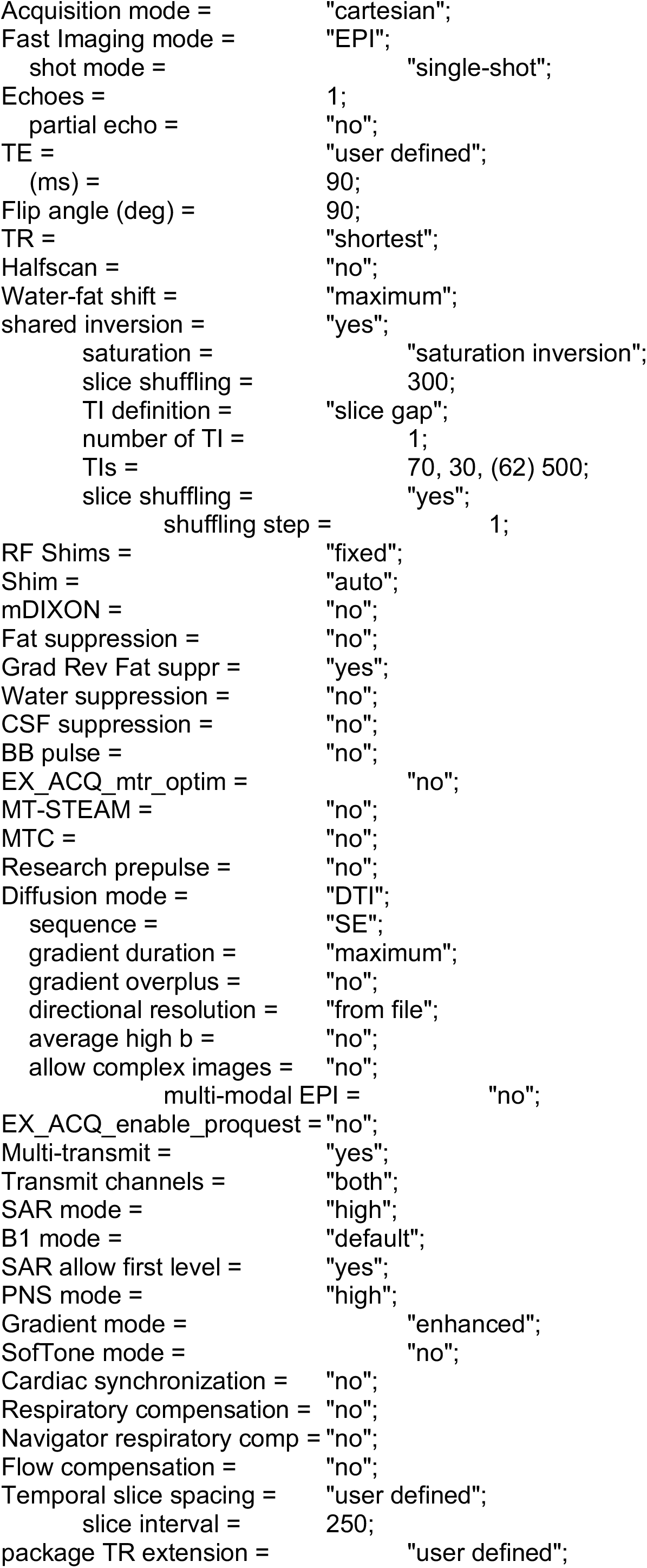
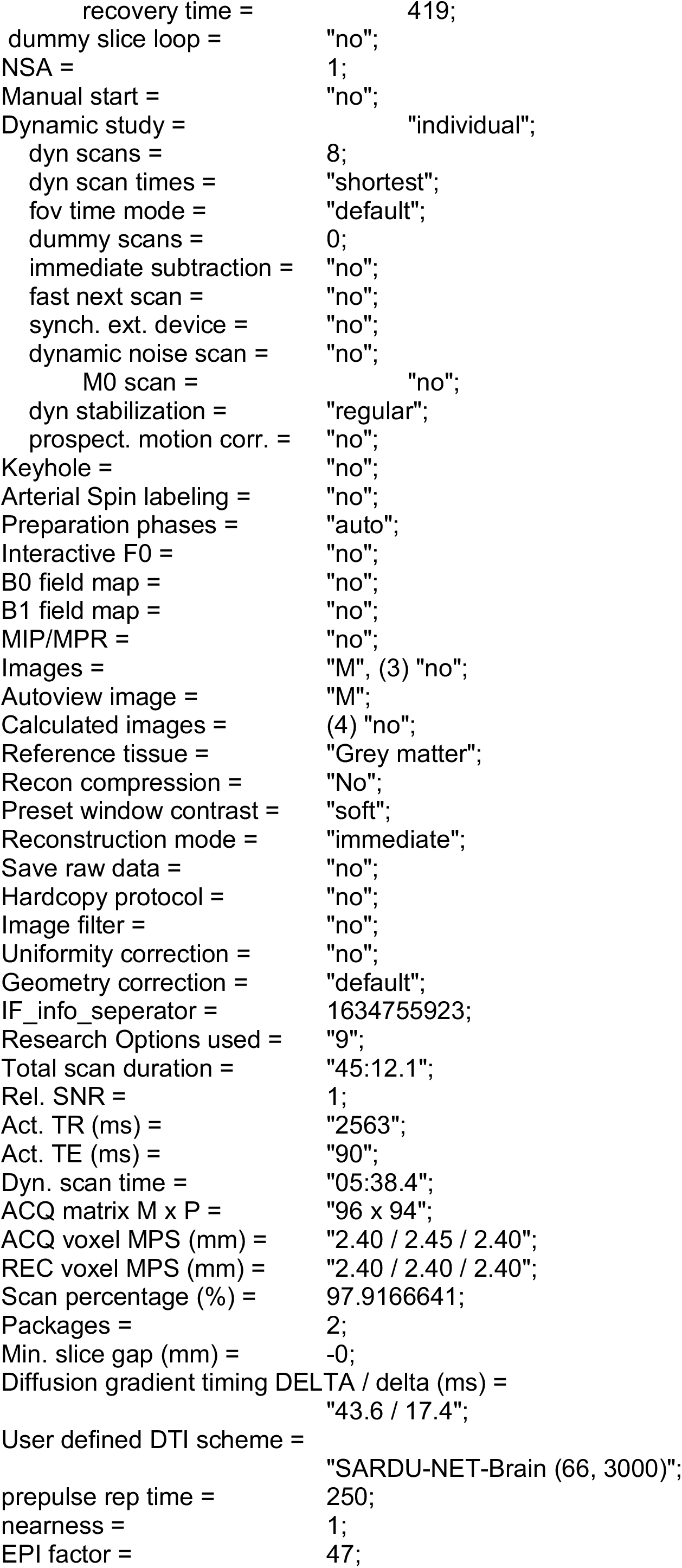
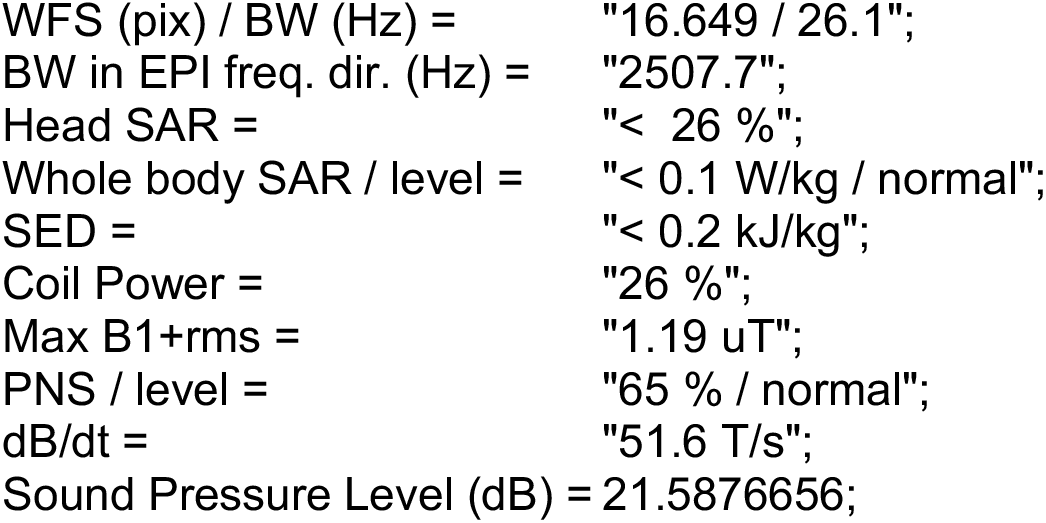
Brain Saturation Inversion Recovery Diffusion imaging, 3T Philips Ingenia CX

**Supporting Information Table S2.**
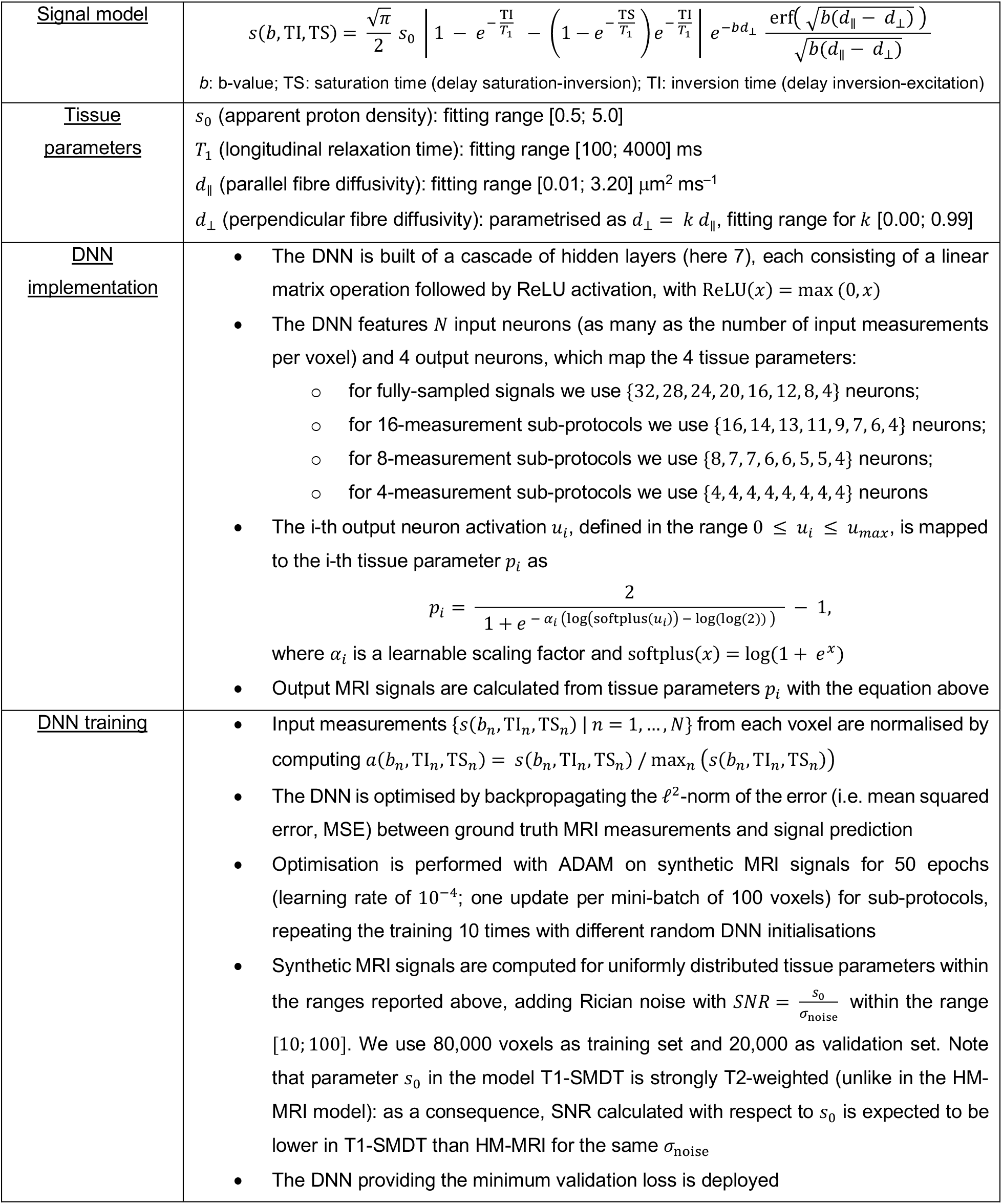
T1-weighted spherical mean diffusion tensor (T1-SMDT) model fitting based on deep neural networks

**Supporting Information Table S3.**
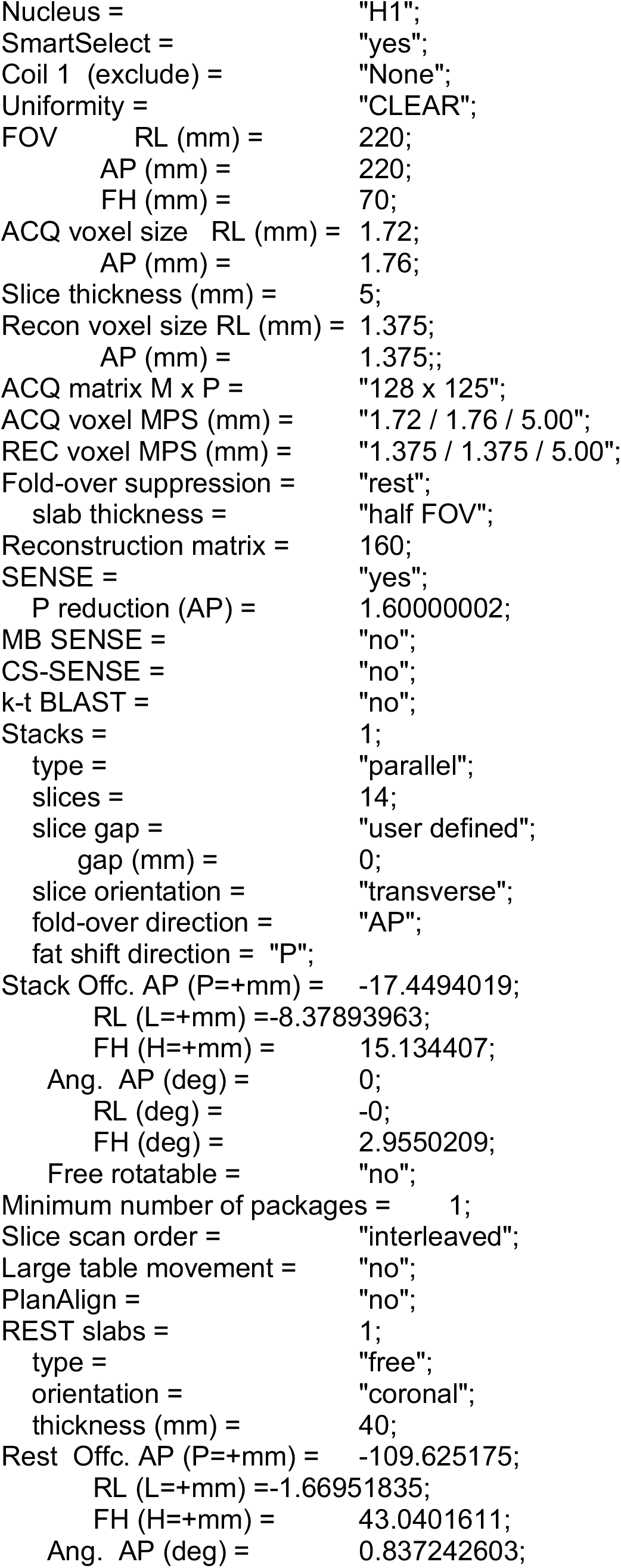
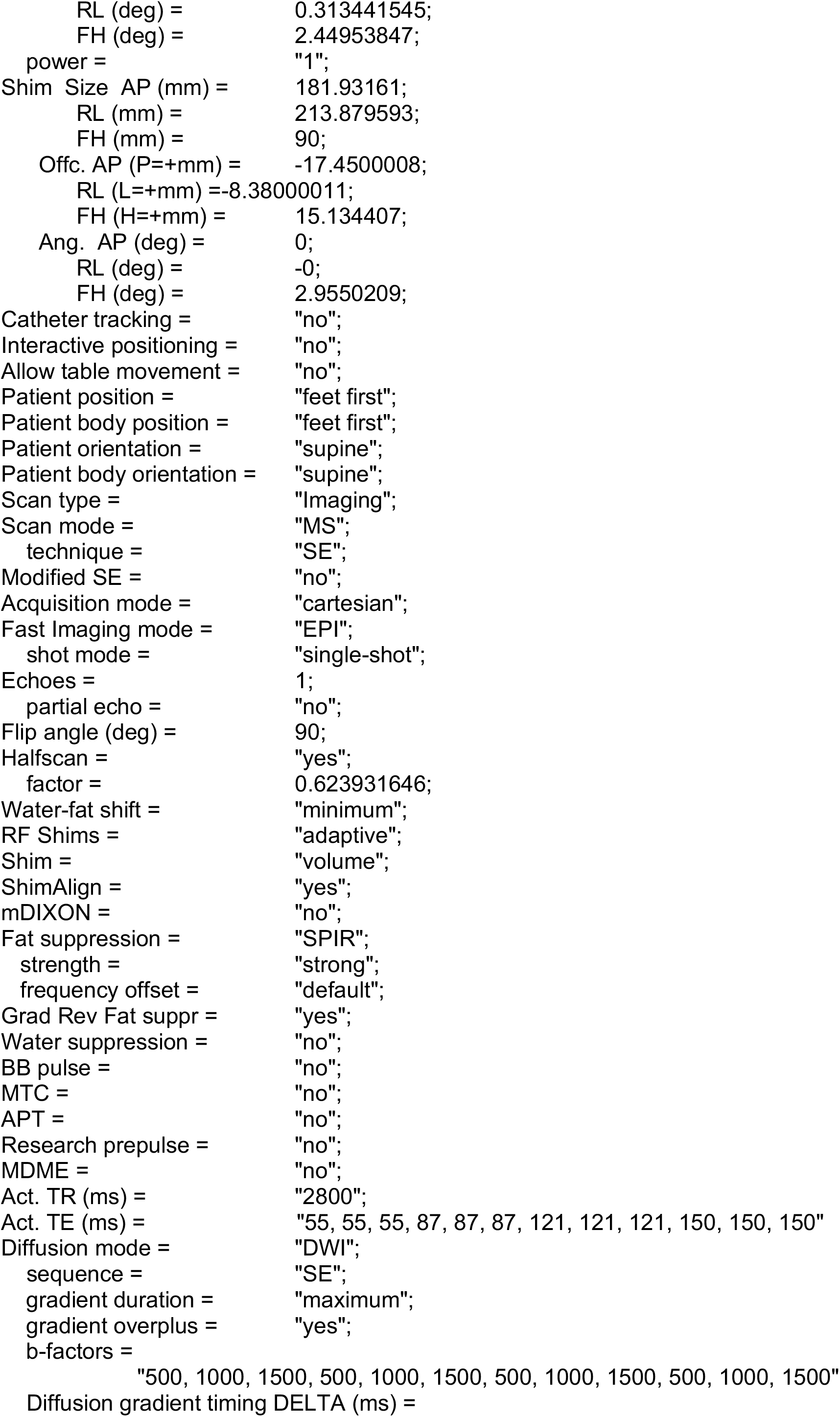
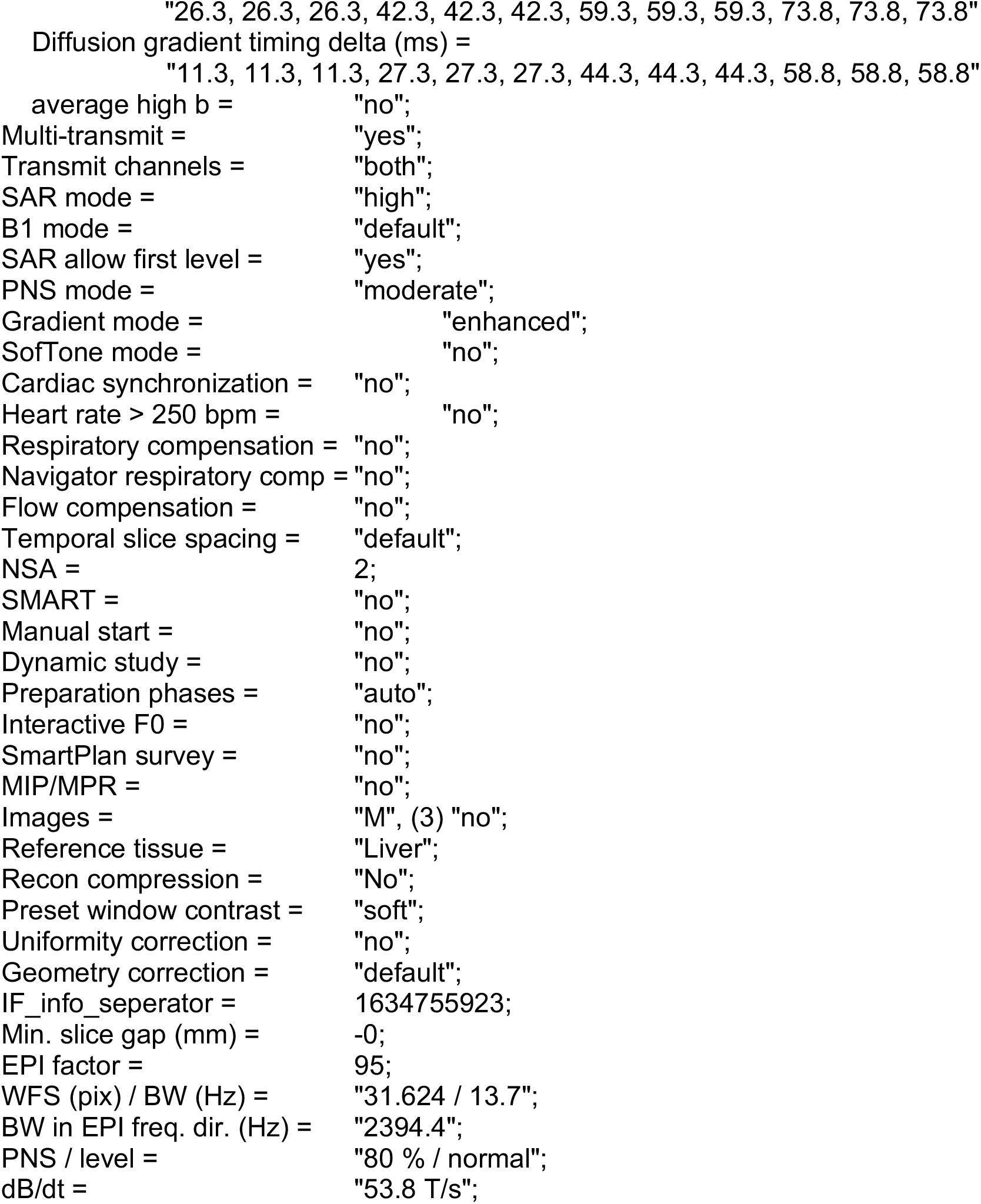
Prostate Diffusion Imaging at varying Echo Time, 3T Philips Achieva

**Supporting Information Table S4.** Hybrid-multidimensional MRI (HM-MRI) model fitting based on deep neural networks

|  |  |
| --- | --- |
| <u>Signal model</u> | $s(b, TE) = s_0 \left( v_l e^{-b d_l - \frac{TE}{T_{2l}}} + (1 - v_l) \left( v_e e^{-b d_e - \frac{TE}{T_{2e}}} + (1 - v_e) e^{-b d_s - \frac{TE}{T_{2s}}} \right) \right)$<br>$b$: b-value; $TE$: echo time |
| <u>Tissue parameters</u> | $s_0$ (apparent proton density): fitting range [0.5; 10.0]<br>$v_l$ (luminal water voxel volume fraction): fitting range [0.0; 1.0]<br>$v_e$ (epithelial fraction of non-luminal tissue): fitting range [0.0; 1.0]<br>$d_l$ (luminal water diffusivity): set to $2.5 \mu\text{m}^2 \text{ms}^{-1}$<br>$d_e$ (epithelial water diffusivity): set to $0.4 \mu\text{m}^2 \text{ms}^{-1}$<br>$d_s$ (stromal water diffusivity): set to $1.4 \mu\text{m}^2 \text{ms}^{-1}$<br>$T_{2l}$ (luminal water T2): set to 650 ms<br>$T_{2e}$ (epithelial water T2): set to 45 ms<br>$T_{2s}$ (stromal water T2): set to 95 ms |
| <u>DNN implementation</u> | The DNN is built of a cascade of hidden layers (here 7), each consisting of a linear matrix operation followed by ReLU activation, with $\text{ReLU}(x) = \max(0, x)$<br>The DNN features $N$ input neurons (as many as the number of input measurements per voxel) and 9 output neurons, which map the 9 tissue parameters:<br>for fully-sampled signals we use {16, 15, 14, 13, 12, 11, 10, 9} neurons;<br>for 12-measurement sub-protocols we use {12, 12, 11, 11, 10, 10, 9, 9} neurons;<br>for 9-measurement sub-protocols we use {9, 9, 9, 9, 9, 9, 9, 9} neurons<br>The $i$-th output neuron activation $u_i$, defined in the range $0 \leq u_i \leq u_{\max}$, is mapped to the $i$-th tissue parameter $p_i$ as $p_i = \frac{2}{1 + e^{-\alpha_i (\log(\text{softplus}(u_i)) - \log(\log(2)))}} - 1,$<br>where $\alpha_i$ is a learnable scaling factor and $\text{softplus}(x) = \log(1 + e^x)$<br>Output MRI signals are calculated from tissue parameters $p_i$ with the equation above |
| <u>DNN training</u> | Input measurements $\{s(b_n, TE_n) \mid n = 1, \dots, N\}$ from each voxel are normalised by computing $a(b_n, TE_n) = s(b_n, TE_n) / \max_n (s(b_n, TE_n))$<br>The DNN is optimised by backpropagating the $\ell^2$-norm of the error (i.e. mean squared error, MSE) between ground truth MRI measurements and signal prediction<br>Optimisation is performed with ADAM on synthetic MRI signals for 50 epochs (learning rate of $10^{-4}$; one update per mini-batch of 100 voxels) for sub-protocols, repeating the training 10 times with different random DNN initialisations<br>Synthetic MRI signals are computed for uniformly distributed tissue parameters within the ranges reported above, adding Rician noise with $SNR = \frac{s_0}{\sigma_{\text{noise}}}$ within the range [35; 250]. We use 80,000 voxels as training set and 20,000 as validation set<br>The DNN providing the minimum validation loss is deployed |

**Supporting Information Table S5.** **TABLE S5** SARDU-Net measurement selection in each of the 3 leave-one-out folds performed on brain DRI. The table shows results for the selection of *D* = 16, *D* = 8 and *D* = 4 measurements out of *M* = 32. In training fold 1, 2 and 3, subject 1, 2 and 3 was respectively left out as test set and not used to train a SARDU-Net via back-propagation. The table also reports the best learning option in each fold (i.e. those providing the lowest validation loss).

|  | Fold 1 | Fold 2 | Fold 3 |
| --- | --- | --- | --- |
| **($b$, TI)**<br>**selection for**<br>**$D = 16$**<br>**[(s mm<sup>-2</sup>, ms)]** | (0,70), (2000,320),<br>(0,570), (0,820),<br>(1000,820), (0,1070),<br>(1000,1070), (3000,1070),<br>(0,1320), (2000,1320),<br>(0,1570), (1000,1570),<br>(2000,1570), (0,1820),<br>(1000,1820), (3000,1820) | (0,70), (3000,320),<br>(0,570), (1000,570),<br>(0,820), (2000,820),<br>(0,1070), (1000,1070),<br>(0,1320), (1000,1320),<br>(0,1570), (1000,1570),<br>(3000,1570), (0,1820),<br>(1000,1820), (3000,1820) | (3000,320), (0,570),<br>(3000,570), (0,820),<br>(2000,820), (0,1070),<br>(1000,1070), (0,1320),<br>(1000,1320), (2000,1320),<br>(0,1570), (1000,1570),<br>(2000,1570), (3000,1570),<br>(0,1820), (1000,1820) |
| | <u>Best learning options:</u><br>100-voxel mini-batch;<br>learning rate of $10^{-3}$;<br>dropout of 0.0 | <u>Best learning options:</u><br>100-voxel mini-batch;<br>learning rate of $10^{-3}$;<br>dropout of 0.0 | <u>Best learning options:</u><br>100-voxel mini-batch;<br>learning rate of $10^{-3}$;<br>dropout of 0.0 |
| **($b$, TI)**<br>**selection for**<br>**$D = 8$**<br>**[(s mm<sup>-2</sup>, ms)]** | (0,820), (3000,820),<br>(0,1070), (0,1320),<br>(2000,1320), (0,1570),<br>(0,1820), (1000,1820) | (2000,320), (0,820),<br>(2000,820), (2000,1070),<br>(0,1320), (0,1570),<br>(0,1820), (1000,1820) | (0,320), (0,820),<br>(2000,1070), (0,1320),<br>(1000,1320), (0,1570),<br>(0,1820), (1000,1820) |
| | <u>Best learning options:</u><br>100-voxel mini-batch;<br>learning rate of $10^{-3}$;<br>dropout of 0.0 | <u>Best learning options:</u><br>1500-voxel mini-batch;<br>learning rate of $10^{-3}$;<br>dropout of 0.0 | <u>Best learning options:</u><br>100-voxel mini-batch;<br>learning rate of $10^{-3}$;<br>dropout of 0.0 |
| **($b$, TI)**<br>**selection for**<br>**for $D = 4$**<br>**[(s mm<sup>-2</sup>, ms)]** | (0,1070), (3000,1570),<br>(0,1820), (1000,1820) | (0,1320), (1000,1320),<br>(0,1820), (3000,1820) | (0,1070), (3000,1320),<br>(0,1820), (1000,1820) |
| | <u>Best learning options:</u><br>100-voxel mini-batch;<br>learning rate of $10^{-4}$;<br>dropout of 0.0 | <u>Best learning options:</u><br>1500-voxel mini-batch;<br>learning rate of $10^{-3}$;<br>dropout of 0.0 | <u>Best learning options:</u><br>100-voxel mini-batch;<br>learning rate of $10^{-3}$;<br>dropout of 0.0 |

**Supporting Information Figure S6.**
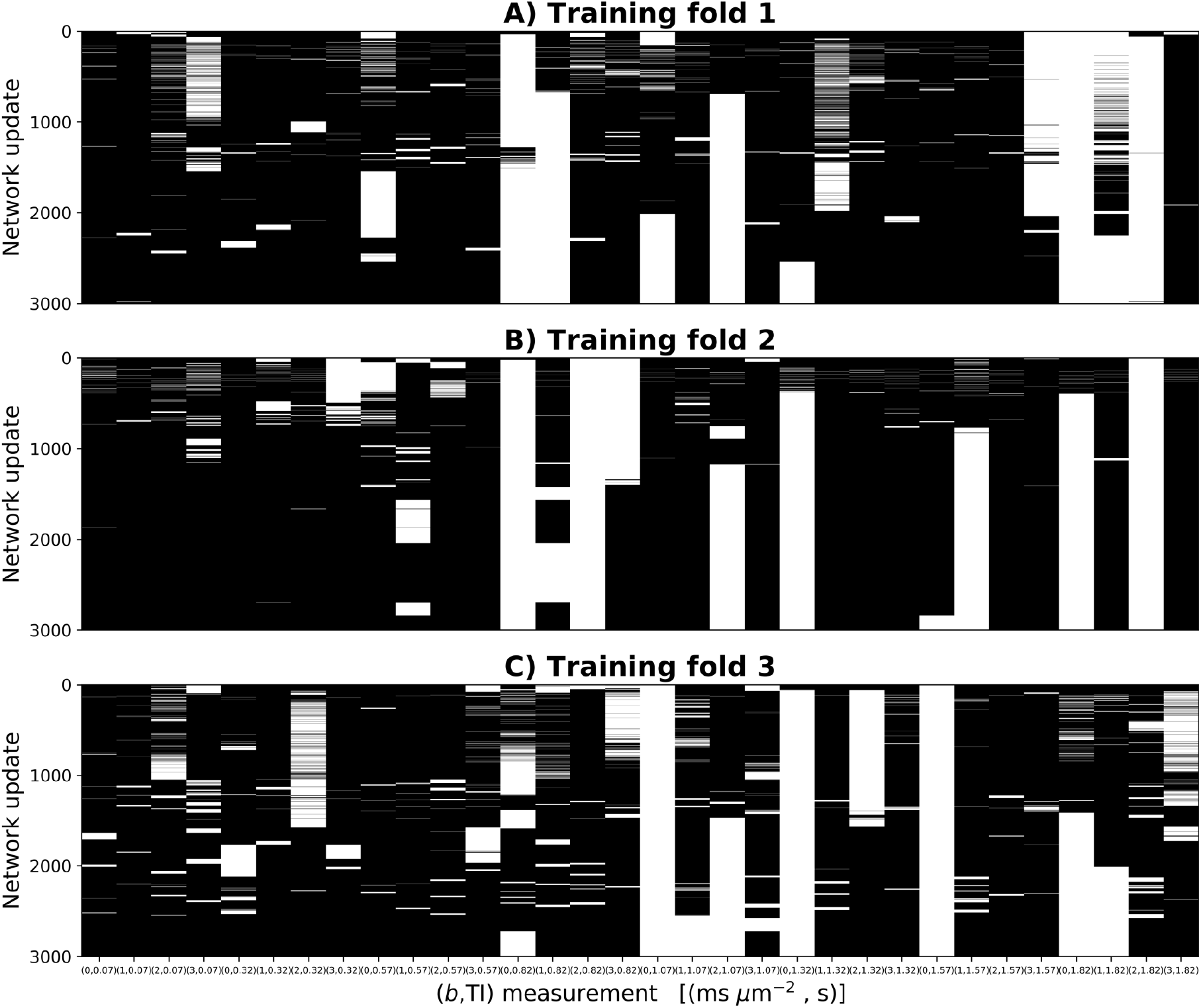
**FIGURE S6** Illustration of measurement selection during SARDU-Net training on *in vivo* brain DRI data. As an example, the figure illustrates the selection of *D* = 8 out of *M* = 32 (*b*,TI) measurements. In each panel, an image illustrates in white measurements that are selected by SARDU-Net as training progresses, while black is used for measurements that are not selected. Different columns refer to different (*b*,TI) measurements, while rows refer to different network parameter updates (here we report the first 3000 updates). Panel A (top), B (middle) and C (bottom) respectively refer to the case when training is performed leaving subject 1, 2 and 3 out as test set (training folds 1, 2 and 3).

**Supporting Information Table S7.** **TABLE S7** SARDU-Net measurement selection in each of the 3 leave-one-out folds performed on prostate DRI. The table shows results for the selection of both *D* = 12 and *D* = 9 measurements out of *M* = 16. In training fold 1, 2 and 3, subject 1, 2 and 3 was respectively left out as test set and not used to train a SARDU-Net via back-propagation. The table also reports the best learning option in each fold (i.e. those providing the lowest validation loss).

|  | Fold 1 | Fold 2 | Fold 3 |
| --- | --- | --- | --- |
| **(b, TE)**<br>**selection for**<br>**$D = 12$**<br>**[(s mm<sup>-2</sup>, ms)]** | (0,55), (500,55),<br>(1000,55), (1500,55),<br>(0,87), (500,87),<br>(1000,87), (1500,87),<br>(0,121), (500,121),<br>(0,150), (500,150) | (0,55), (500,55),<br>(1000,55), (1500,55),<br>(0,87), (500,87),<br>(1000,87), (1500,87),<br>(0,121), (500,121),<br>(0,150), (500,150) | (0,55), (500,55),<br>(1000,55), (1500,55),<br>(0,87), (500,87),<br>(1000,87), (1500,87),<br>(0,121), (0,150),<br>(500,150), (1000,150) |
|  | <u>Best learning options:</u><br>100-voxel mini-batch;<br>learning rate of 10<sup>-3</sup>;<br>dropout of 0.0 | <u>Best learning options:</u><br>100-voxel mini-batch;<br>learning rate of 10<sup>-3</sup>;<br>dropout of 0.0 | <u>Best learning options:</u><br>100-voxel mini-batch;<br>learning rate of 10<sup>-3</sup>;<br>dropout of 0.0 |
| **(b, TE)**<br>**selection for**<br>**$D = 9$**<br>**[(s mm<sup>-2</sup>, ms)]** | (0,55), (500,55),<br>(1000,55), (0,87),<br>(1500,87), (0,121),<br>(500,121), (0,150),<br>(500,150) | (0,55), (500,55),<br>(1000,55), (0,87),<br>(500,87), (1000,87),<br>(500,121), (1000,121),<br>(0,150) | (0,55), (1000,55),<br>(1500,55), (0,87),<br>(500,87), (1000,87),<br>(500,121), (0,150),<br>(500,150) |
|  | <u>Best learning options:</u><br>100-voxel mini-batch;<br>learning rate of 10<sup>-3</sup>;<br>dropout of 0.0 | <u>Best learning options:</u><br>100-voxel mini-batch;<br>learning rate of 10<sup>-3</sup>;<br>dropout of 0.0 | <u>Best learning options:</u><br>100-voxel mini-batch;<br>learning rate of 10<sup>-3</sup>;<br>dropout of 0.0 |

**Supporting Information Table S8.**
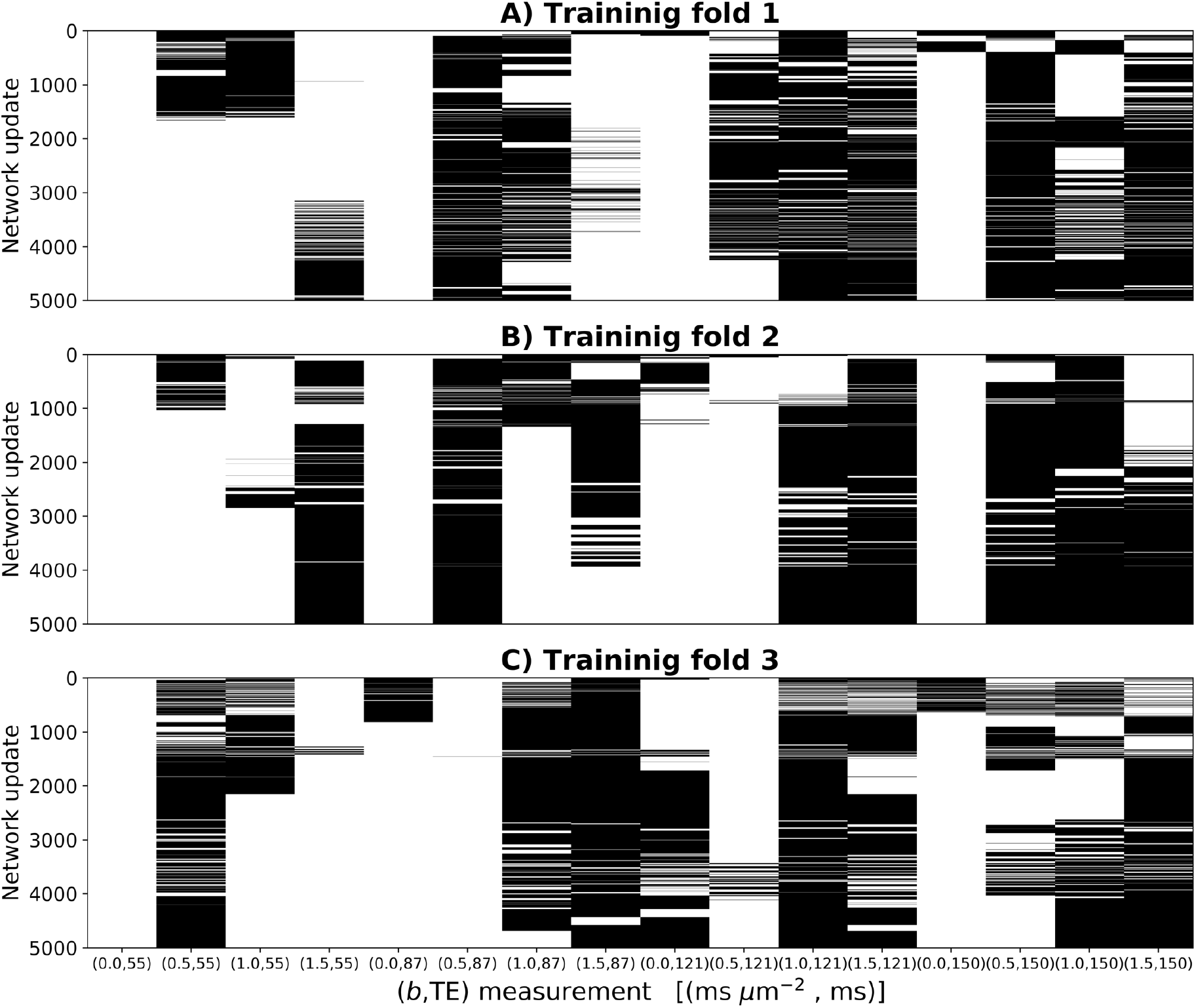
**FIGURE S8** Illustration of measurement selection during SARDU-Net training on *in vivo* prostate DRI data. As an example, the figure illustrates the selection of *D* = 9 out of *M* = 16 (*b*,TE) measurements. In each panel, an image illustrates in white measurements that are selected by SARDU-Net as training progresses, while black is used for measurements that are not selected. Different columns refer to different (*b*,TE) measurements, while rows refer to different network parameter updates (here we report the first 5000 updates). Panel A (top), B (middle) and C (bottom) respectively refer to the case when training is performed leaving subject 1, 2 and 3 out as test set (training folds 1, 2 and 3).

## Notes

https://github.com/fragrussu/sardunet

### Competing Interest Statement

Francesco Grussu is supported by PREdICT, a study co-funded by AstraZeneca in Spain. Torben Schneider is an employee of Deep Spin (Germany) and previously worked for Philips (UK).

### Summary of Updates

This version of the manuscript has been revised to include a demonstration of SARDU-Net for protocol design in diffusion-relaxation imaging of the brain. The new data consist of saturation inversion recovery diffusion-weighted imaging performed on 3 healthy volunteers at 3. A rich diffusion-/T1-weighting protocol made of 32 unique b-values and inversion times.

